# Missense mutations in the MLKL ‘brace’ region lead to lethal neonatal inflammation in mice and are present in high frequency in humans

**DOI:** 10.1101/628370

**Authors:** Joanne M. Hildebrand, Maria Kauppi, Ian J. Majewski, Zikou Liu, Allison Cox, Sanae Miyake, Emma J. Petrie, Michael A. Silk, Zhixiu Li, Maria C. Tanzer, Samuel N. Young, Cathrine Hall, Sarah E. Garnish, Jason Corbin, Michael D. Stutz, Pradnya Gangatirkar, Emma C. Josefsson, Kristin Rigbye, Holly Anderton, James A. Rickard, Anne Tripaydonis, Julie Sheridan, Thomas S. Scerri, Peter A. Czabotar, Jian-Guo Zhang, Cody C. Allison, Marc Pellegrini, Gillian M. Tannahill, Esme C. Hatchell, Tracy A. Willson, Dina Stockwell, Carolyn A. de Graaf, Janelle Collinge, Adrienne Hilton, Natasha Silke, Sukhdeep K. Spall, Diep Chau, Vicki Athanasopoulos, Donald Metcalf, Ronald M. Laxer, Alexander G. Bassuk, Benjamin W. Darbro, Maria A. Fiatarone Singh, Nicole Vlahovich, David Hughes, Maria Kozlovskaia, David B. Ascher, Klaus Warnatz, Nils Venhoff, Jens Thiel, Stefan Blum, John Reveille, Michael S. Hildebrand, Carola G. Vinuesa, Pamela McCombe, Matthew A. Brown, Ben T. Kile, Catriona McLean, Melanie Bahlo, Seth L. Masters, Hiroyasu Nakano, Polly J. Ferguson, James M. Murphy, Warren S. Alexander, John Silke

## Abstract

We have isolated a mouse strain with a single missense mutation in the gene encoding MLKL, the essential effector of necroptotic cell death. The resulting substitution lies within the two-helix ‘brace’ and confers constitutive, RIPK3 independent, killing activity to MLKL. Mice homozygous for *Mlkl^D139V^* develop lethal inflammation within days of birth, implicating the salivary glands and pericardium as hotspots for necroptosis and inflammatory infiltration. The normal development of *Mlkl^D139V^* homozygotes until birth, and the absence of any overt phenotype in heterozygotes provides important *in vivo* precedent for the capacity of cells to clear activated MLKL. These observations offer an important insight into the potential disease-modulating roles of three common human *MLKL* polymorphisms that encode amino acid substitutions within or adjacent to the brace region. Compound heterozygosity of these variants is found at up to 12-fold the expected frequency in patients that suffer from a pediatric autoinflammatory disease, CRMO.

## INTRODUCTION

Necroptosis is a form of programmed cell death associated with the production of pro-inflammatory cytokines, destruction of biological membranes and the release of intracellular Damage Associated Molecular Patterns (DAMPs) (Newton and Manning, 2016). Necroptosis depends on the activation of pseudokinase Mixed Lineage Kinase domain-Like (MLKL) by Receptor Interacting Protein Kinase 3 (RIPK3) (Murphy et al., 2013; Sun et al., 2012; Zhao et al., 2012). RIPK3-mediated phosphorylation of MLKL triggers a conformational change that facilitates the translocation to, and eventual irreversible disruption of, cellular membranes. While the precise biophysical mechanism of membrane disruption is still a matter of debate, it is consistently associated with the formation of an MLKL oligomer and the direct association of the four-helix bundle domain (4HB) of MLKL with membranes (Cai et al., 2014; Chen et al., 2014; Dondelinger et al., 2014; Hildebrand et al., 2014). In mouse cells, the expression of the murine MLKL 4HB domain alone (residues 1-125), 4HB plus brace helix (1-180), or the expression of phosphomimetic or other single site pseudokinase domain (PsKD) mutants is sufficient to induce membrane translocation, oligomerization and membrane destruction (Hildebrand et al., 2014; Murphy et al., 2013). While capable of disrupting synthetic liposomes when produced recombinantly, similarly truncated and equivalent single site (PsKD) mutant forms of human MLKL do not robustly induce membrane associated oligomerization and cell death without forced dimerization (Petrie et al., 2018; Quarato et al., 2016; Tanzer et al., 2016). Furthermore, both mouse and human MLKL mutants have been reported that have the capacity to form membrane associated oligomers, but fail to cause irreversible membrane disruption and cell death (Hildebrand et al., 2014; Petrie et al., 2018). Recent studies have revealed that necroptosis downstream of MLKL phosphorylation and membrane association can be modulated by processes that utilize the Endosomal Sorting Complex Required for Transport (ESCRT) family of proteins. One model proposes a role for ESCRT in limiting necroptosis via plasma membrane excision and repair (Gong et al., 2017) while other models limit plasma membrane disruption by ESCRT-mediated endosomal trafficking and the release of MLKL in endosomes (Yoon et al., 2017) or the shedding of phosphorylated MLKL in extracellular vesicles (Zargarian et al., 2017).

In mice, the absence of MLKL does not appear to have obvious deleterious developmental or homeostatic effects (Murphy et al., 2013; Wu et al., 2013). However, genetic deletion of *Fadd*, *Casp8* or *Ripk1*, leads to inappropriate activation of MLKL and ensuing necroptosis during embryogenesis and is incompatible with life beyond embryonic day (E)10.5, E10.5 and 1-3 days post-natally, respectively (Dillon et al., 2014; Kaiser et al., 2014; Kelliher et al., 1998; Rickard et al., 2014b; Varfolomeev et al., 1998; Yeh et al., 1998; Zhang et al., 2011). Exploring the precise physiological consequences of inappropriate MLKL activation in these scenarios is complicated by the fact that FADD, Caspase-8 and RIPK1 also play important roles in cellular processes other than modulation of MLKL-induced necroptotic cell death (Alvarez-Diaz et al., 2016; Kaiser et al., 2011; Kang et al., 2004; Newton et al., 1998; Oberst et al., 2011; Rickard et al., 2014b).

Aberrant levels of MLKL-dependent cell death contribute to disease in several genetic and experimental mouse models (Anderton et al., 2017; Dannappel et al., 2014; Hockendorf et al., 2016; Newton et al., 2016; Rickard et al., 2014a; Rickard et al., 2014b). In humans, *MLKL* mRNA and protein levels are positively correlated with survival of patients with pancreatic adenocarcinoma, cervical-, gastric-, ovarian- and colon-cancers (reviewed by (Lalaoui and Brumatti, 2017)). Interestingly, high levels of phosphorylated MLKL are associated with reduced survival in esophageal and colon cancer patients (Liu et al., 2016b). Two missense *MLKL* somatic mutations identified in human cancer tissue have been found to confer a reduction in necroptotic function in cell-based assays (Murphy et al., 2013; Petrie et al., 2018). One recent study reported a significant enrichment of an ultra rare *MLKL* stop-gain gene variant p.Q48X in Hong Kong Chinese patients suffering from a form of Alzheimer’s disease (Wang et al., 2018) however more common germline *MLKL* gene variants are only weakly associated with human disease in GWAS databases. In two recent studies, lethal immunodeficiency, arthritis and intestinal inflammation was reported in patients homozygous for ultra rare-loss of function RIPK1 mutations (Cuchet-Lourenco et al., 2018; Li et al., 2019), however to date, *MLKL* gene variants have not been directly implicated in any severe Mendelian forms of human disease.

We have identified a single base pair germline mutation of mouse *Mlkl* that encodes a missense alteration to the MLKL mouse brace region and confers constitutive activation independent of upstream necroptotic stimuli. Given this mutant *Mlkl* allele is subject to the same developmental and environmental controls on gene expression as wildtype *Mlkl*, the postnatal lethality in these mice provides novel insight into the physiological and pathological consequences of dysregulated necroptosis. In parallel these findings inform the potential functional significance of three common human *MLKL* polymorphisms that encode non-conservative amino acid substitutions within, or in close proximity to, the brace helix that is mutated in the *Mlkl^D139V^* mouse.

## RESULTS

### Generation of a constitutively active form of MLKL

An ENU mutagenesis screen was performed to identify mutations that ameliorate thrombocytopenia in *Mpl^−/−^* mice (Kauppi et al., 2008). A G_1_ founder, designated *Plt15*, had a modestly elevated platelet count of 189×10^6^/mL compared to the mean for *Mpl^−/−^* animals (113±57×10^6^/mL) and yielded 19 *Mpl^−/−^* progeny. Ten of these mice had platelet counts over 200×10^6^/mL, consistent with segregation of a dominantly acting mutation (Fig. 1A). Linkage analysis and sequencing (see Experimental Procedures) identified an A to T transversion in *Mlkl* that was heterozygous in all mice with an elevated platelet count (Fig. 1B). The *Mlkl^Plt15^* mutation results in a non-conservative aspartic acid-to-valine substitution at position 139. In the full length mMLKL structure D139 forms a salt bridge with an arginine residue at position 30 (α2 helix) of the MLKL four-helix bundle (4HB) domain (Murphy et al, 2013) (Fig. 1C). This salt bridge represents one of a series of electrostatic interactions between residues in helix α2 of the MLKL 4HB domain and the two-helix ‘brace’ region. D139 of mouse MLKL is conserved in all MLKL orthologues in vertebrata reported to date (Fig. 1D). We have shown that the exogenous expression of the 4HB domain of murine MLKL alone is sufficient to kill mouse fibroblasts whereas exogenous expression of full length MLKL does not, indicating that this ‘electrostatic zipper’ may play an important role in suppressing the killing activity of the MLKL 4HB (Hildebrand et al., 2014). To determine if MLKL^D139V^ exhibited altered ability to induce necroptotic cell death relative to MLKL^Wt^, we stably expressed these full length proteins under the control of a doxycycline-inducible promoter in immortalized mouse dermal fibroblasts (MDF) isolated from *Wt, Mlkl^−/−^, Ripk3^−/−^* or *Ripk3^−/−^;Casp8^−/−^* mice. While expressed at comparable levels, MLKL^D139V^ induced markedly more death than MLKL^Wt^, on each of the genetic backgrounds tested (Fig. 1E-F, Supp. Fig. 1A). This indicates that MLKL^D139V^ is a constitutively active form of MLKL, capable of inducing necroptotic cell death independent of upstream signaling and phosphorylation by its activator RIPK3. Consistent with this interpretation, exogenous expression of MLKL^D139V^ in *Ripk3^−/−^;Casp8^−/−^* MDFs was sufficient to induce the organelle swelling and plasma membrane rupture characteristic of TNF induced necroptosis when examined by Transmission Electron Microscopy (Fig. 1G).

**Figure 1:**
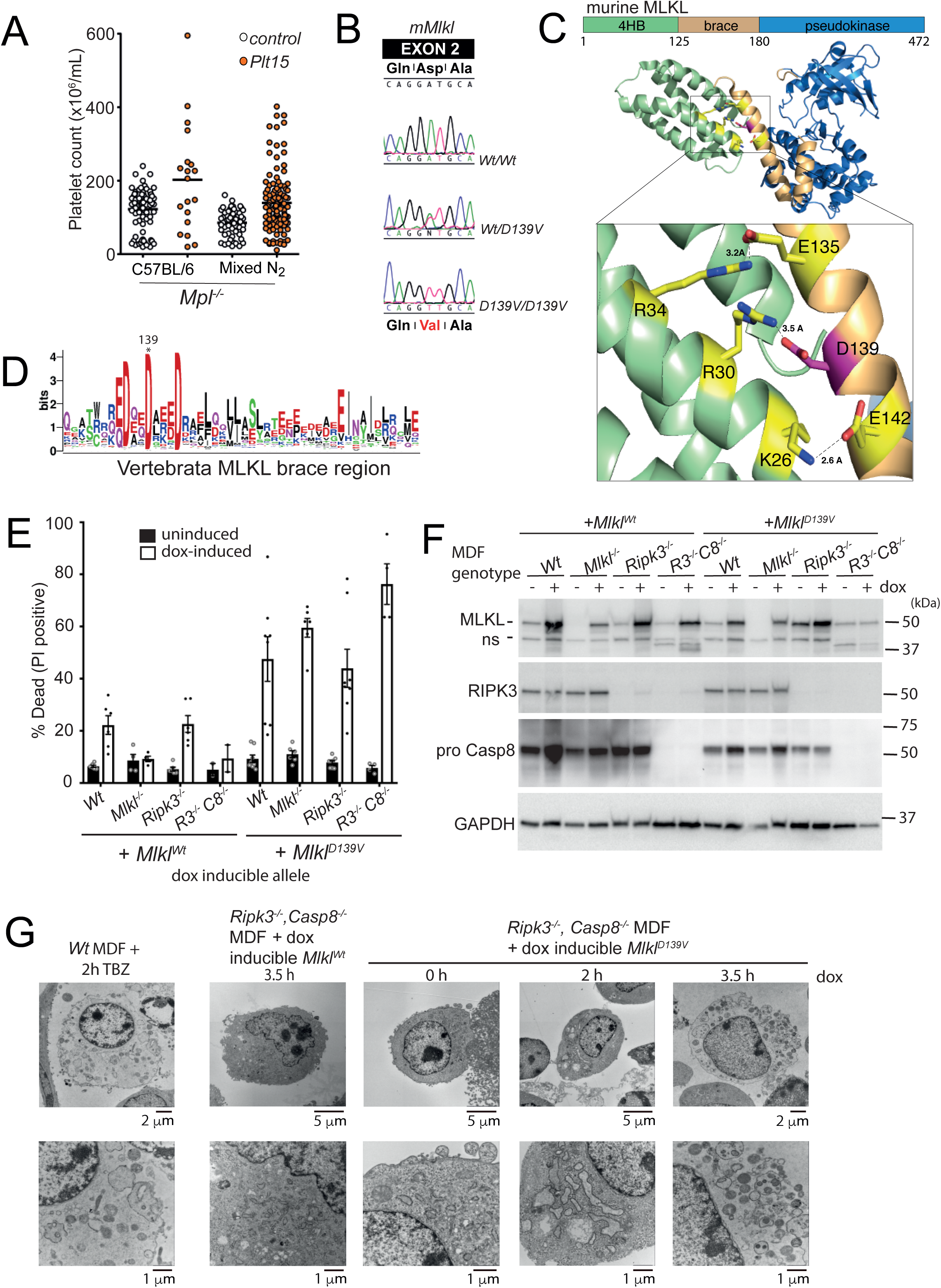
Murine MLKL^D139V^ is a constitutively active form of MLKL. (A) Platelet counts from *Mpl^−/−^* mice and offspring from matings between *Plt15* mice and *Mpl^−/−^* mice on a C57BL/6 or mixed C57BL/6:129/Sv background used for linkage analysis (Mixed N_2_). (B) A missense mutation (D139V) in the second exon of *Mlkl* was identified in DNA isolated from *Plt15* mutant mice. DNA sequence is shown for a wild type (top), a heterozygous mutant (middle), and a homozygous mutant (bottom). (C) Aspartate 139 contributes to an ‘electrostatic zipper’ joining brace helix 1 and the 4HB a2 helix of mouse MLKL (PDB code 4BTF) (Murphy et al., 2013). (D) Sequence logo of the MLKL brace domain generated from a multiple sequence alignment of all Vertebrata MLKL sequences (257) available on OrthoDB. (E) Mouse dermal fibroblasts (MDFs) of indicated genotypes were stably transduced with *Mlkl^Wt^* and *Mlkl^D139V^*lentiviral constructs and expression was induced with doxycycline (dox) for 21 hrs. PI-positive cells were quantified by flow cytometry. Means ± SEM are plotted for between 4-8 experiments (a combination of biological repeats and independent experiments) for each genotype with the exception of *R3*^−/−^*C8*^−/−^ + *Mlkl^Wt^* (n=2, ± SD). (F) Western blot analysis of whole cell lysates taken 6 hours post doxycycline induction for analysis of MLKL, RIPK3 and pro-caspase 8 expression. (G) Transmission electron micrographs of MDFs stimulated as indicated. Images selected are representative of 2-3 independent analyses. TBZ; TNF + Birinapant + Z-VAD-FMK.

### Constitutively active mouse MLKL causes a lethal perinatal inflammatory syndrome

To define the phenotypic consequences of constitutively active MLKL in the absence of any confounding effects resulting from *Mpl*-deficiency, all subsequent studies were performed on a *Mpl^+/+^* background. Homozygous *Mlkl^D139V/D139V^* pups were born at expected Mendelian frequencies (Supp. Table I) and were ostensibly normal macroscopically and histologically at E19.5 (Supp. Fig. 2A-D). However, by 3 days of age, although outwardly indistinguishable from littermates (Fig. 2A), they exhibited reduced body weight (Supp. Fig. 2B) and failed to thrive, with a maximum observed lifespan of 6 days under conventional clean housing conditions. Like *Mlkl^Wt/D139V^* mice, *Mlkl^null/D139V^* compound heterozygotes were present at the expected frequency at P21 and developed normally to adulthood (Supp. Table II). Thus, the constitutive activity of MLKL^D139V^ was not affected by the presence of normal MLKL protein suggesting it is the absolute allelic dose of *Mlkl^D139V^* that determines perinatal lethality. To confirm that the phenotype of the ENU derived *Mlkl^D139V^* mice was due to the *Mlkl^D139V^* missense mutation, we independently generated *Mlkl^D139V^* mice using CRISPR-Cas9 genomic editing. Homozygote CRISPR-*Mlkl^D139V/D139V^* mice also died soon after birth (Supp. Table III).

**Figure 2:**
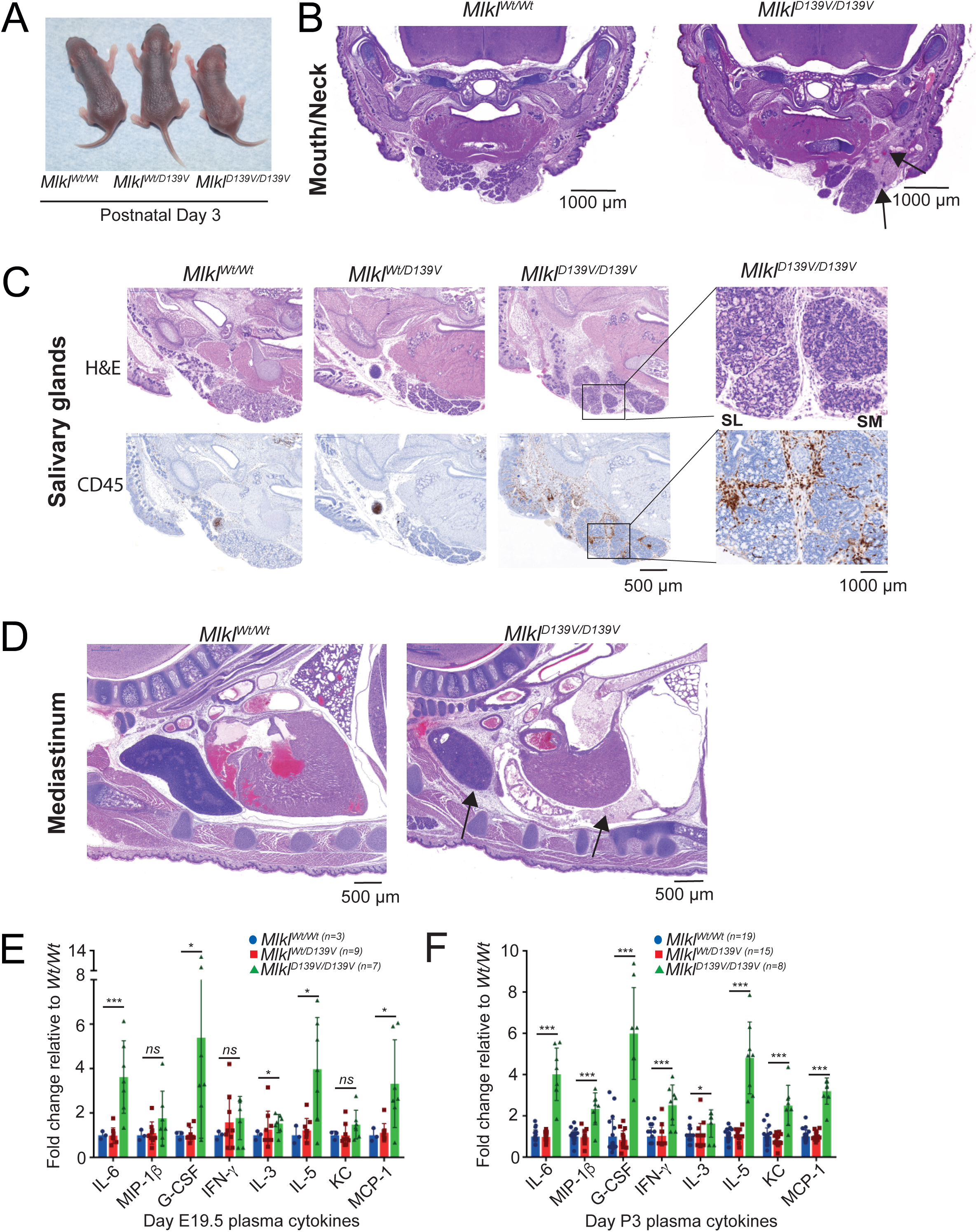
Homozygous *Mlkl^D139V^* neonates exhibit dispersed inflammation and secondary lymphoid organ hypoplasia throughout the head, neck and mediastinum. (A) Macroscopic appearance of *Mlkl^Wt/Wt^*, *Mlkl^Wt/D139V^* and *Mlkl^D139V/D139V^* mice at postnatal day 3. (B) Coronal section of mouth and neck region of postnatal day 2 litter mates stained with haematoxylin and eosin (H&E). Dilated blood vessels and edema are indicated by arrows. (C) Serial mandible sections from postnatal day 3 litter mates stained with H&E and anti-CD45. Inset black boxes are magnified in right panel. SL, sublingual gland. SM, submandibular gland. Images representative of n=3-4 P3 pups per genotype. (D) H&E stained sections from mediastinum of postnatal day 2 litter mates. Thymic cortical thinning and pericardial infiltration are indicated by arrows. For full anatomical annotations for B and D see Supp. Fig. 2H. (B) and (D) representative of n= 5-6 P2 pups examined. Multiplex measurement of plasma cytokines levels at E19.5 (E) and postnatal day 3 (F). Error bars represent mean ± SD of indicated numbers of independent pups per genotype sampled.

Hematoxylin-Eosin stained-sections from both P2 and P3 *Mlkl^D139V/D139V^* pups revealed multifocal acute inflammation characterized by neutrophilic infiltration, dilated blood vessels and edema (Fig. 2B) in the dermis and subcutis of the head and neck. These inflammatory features were not observed in *Mlkl^Wt/Wt^* or *Mlkl^Wt/D139V^* littermates, nor in *Mlkl*^−/−^ mice of the same age (Supp. Fig. 2I). Cells of hematopoietic origin, revealed by immunohistochemical staining for CD45, were sparsely distributed throughout the lower head and neck and confined predominantly to a clearly delineated developing lymph node in *Mlkl^Wt/Wt^* and *Mlkl^Wt/D139V^* littermates (Fig. 2C). In contrast, CD45^+^ cells were more numerous and distributed throughout the cutis, subcutis and salivary glands of *Mlkl^D139V/D139V^* pups (Fig. 2C). A mixture of diffuse and focal inflammatory infiltration was also observed within the mediastinum and pericardial space of all P2/P3 *Mlkl^D139V/D139V^* pups examined, as was a marked paucity of thymic cortical lymphocytes (Fig. 2D, Supp. Fig 2E), phenotypes not evident in E19.5 embryos (Supp. Fig. 2D). Apart from small foci of hepatocyte and enterocyte loss/necrosis evident in the livers and small intestines of some *Mlkl^D139V/D139V^* pups examined (data not shown), no other lesions were observed by histopathology. Consistent with this inflammatory phenotype significantly elevated levels of several pro-inflammatory cytokines and chemokines were evident in the plasma of both E19.5 and P3 *Mlkl^D139V/D139V^* pups (Fig. 2E, F). Blood glucose levels were normal (Supp. Fig. 2 F, G).

### Hematopoetic defects in *Mlkl^D139V^* mice

Although blood cell numbers were unchanged in *Mlkl^D139V/D139V^* pups at E19.5 relative to *Mlkl^Wt/Wt^* and *Mlkl^Wt/D139V^* littermates, by P3 significant deficits were evident in total white blood cell count, lymphocyte and platelet numbers (Fig. 3A-C, Supp. Fig. 3A). Similarly, the numbers of hematopoietic stem and progenitor cells were present at normal proportions in fetal livers of E18.5 *Mlkl^D139V/D139V^* pups, although increased levels of intracellular ROS were uniformly evident (Fig. 3D-E, Supp. Fig. 3B). By P2, deficits in CD150^+^CD48^+^ and CD150^+^CD48^−^ populations were present (Fig. 3F), accompanied by increased AnnexinV binding (which indicates either phosphatidyl serine exposure or plasma membrane rupture) in all lineages (Fig. 3G). In adult *Mlkl^Wt/D139V^* mice, numbers of hematopoietic stem and progenitor cells were unaffected (Fig. 3H); however, upon myelosuppressive irradiation, recovery of hematopoietic cell numbers was delayed and characterized by increased expression of ROS and Annexin V (Supp. Fig. 3C, D). When challenged with the cytotoxic drug 5-fluorouracil (5-FU), blood cell recovery in *Mlkl^Wt/D139V^* mice was similarly delayed (Fig. 3I). In competitive transplants in which test *Mlkl^Wt/D139V^* or *Mlkl^Wt/Wt^* marrow was co-injected with wild type competitor marrow in 10:1 excess, as expected, *Mlkl^Wt/Wt^* marrow contributed to 90% of recipient blood cells 8 weeks after transplantation and maintained that level of contribution for 6 months (Fig. 3J). In contrast, *Mlkl^Wt/D139V^* marrow performed poorly, contributing to 25% and 51% of recipient blood cells at these times (Fig. 3J). Similarly, while wild type fetal liver cells contributed to the vast majority of blood cells in irradiated recipients up to 6 months after transplantation, cells from *Mlkl^D139V/D139V^* embryos failed to compete effectively during this period (Fig. 3K). Heterozygote *Mlkl^Wt/D139V^* fetal liver cells contributed poorly in the first month following the graft but recovered to contribute more after six months (Fig. 3K). Thus, while tolerated under steady-state conditions, heterozygosity of *Mlkl^D139V^* is deleterious under conditions of hematopoietic stress. Bone marrow-derived HSCs from *Mlkl^Wt/D139V^* adults and fetal liver-derived HSCs from *Mlkl^Wt/D139V and^ Mlkl^D139V/D139V^* pups also formed fewer and smaller colonies in the spleens of lethally irradiated recipient mice after 8 days (Supp. Fig. 3E).

**Figure 3:**
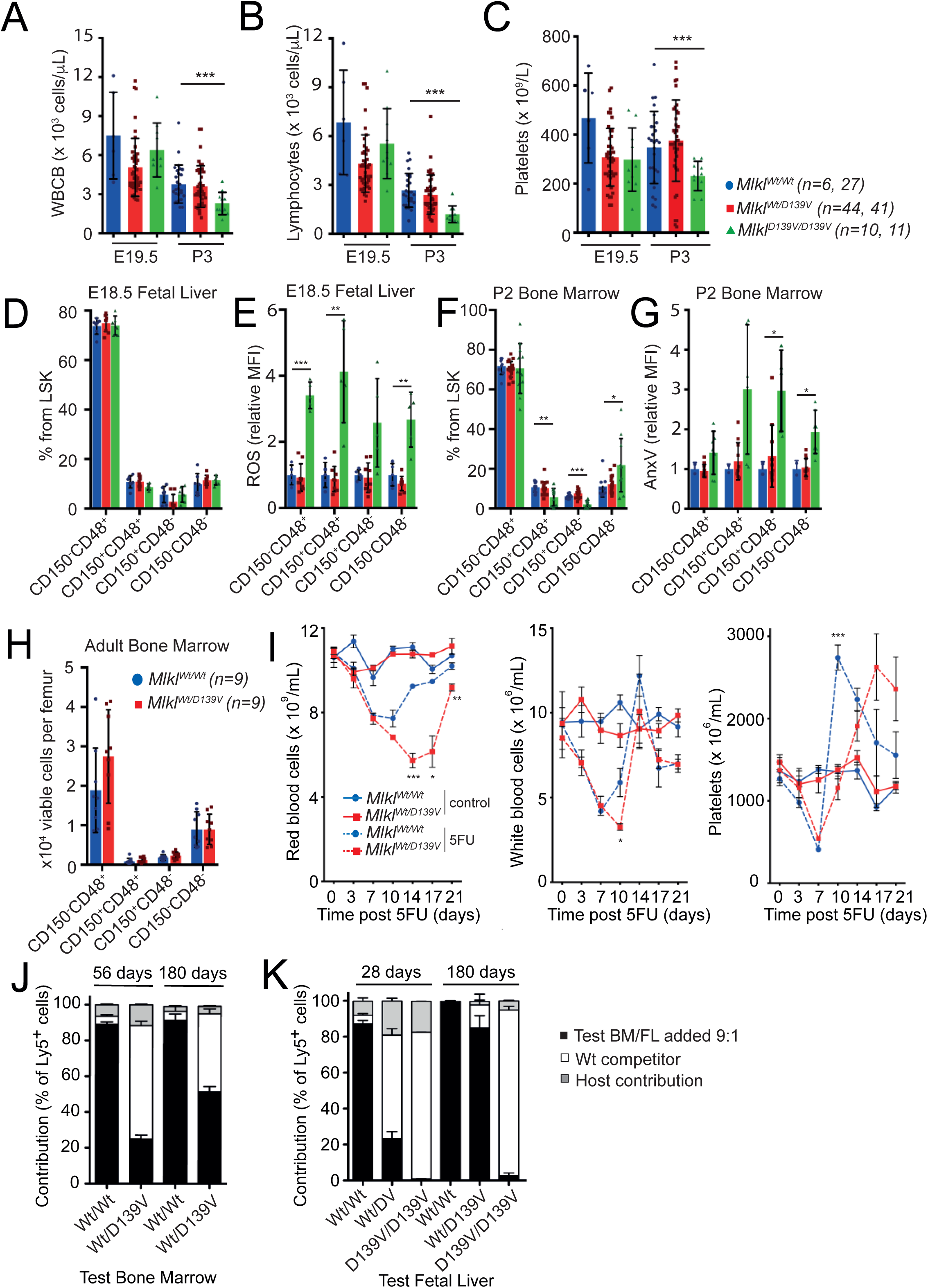
Alterations in hematopoietic cells and defective emergency hematopoiesis in *Mlkl*^D139V^ mice. (A-C) Absolute white blood cell (WBCB) and lymphocyte numbers in the peripheral blood of E19.5 and P3 pups, n indicated. (D-G) Proportions of HSC (Lineage^−^Sca-1^+^c-kit^+^ (LSK) CD150^+^ CD48^−^), MPP (LSK CD150^−^ CD48^−^), HPC-1 (LSK CD150^−^ CD48^+^) and HPC-2 (LSK CD150^+^ CD48^+^)(Oguro et al., 2013) in E18.5 fetal liver (D) and P2 bone marrow cells (F). Levels of ROS in E18.5 fetal liver cell populations (E) and AnnexinV expression in P2 bone marrow cell populations (G) are shown relative to *Mlkl^Wt^/^Wt^*. *Mlkl^Wt/Wt^* -blue bar, Mlkl*^Wt/D139V^* ^−^red bar, MlklD^139V/D139V-^green bar. Values from all independent animals sampled are plotted (n=2-18). (H) Numbers of HSC (Lineage^−^Sca-1^+^c-kit^+^ (LSK) CD150^+^ CD48^−^), MPP (LSK CD150^−^ CD48^−^), HPC-1 (LSK CD150^−^ CD48^+^) and HPC-2 (LSK CD150^+^ CD48^+^) in adult bone marrow from *Mlkl^Wt/Wt^* and *Mlkl^Wt/D139V^* mice, n indicated. Error bars in A-G represent mean ± SD. (I) Numbers of red and white blood cells and platelets in *Mlkl^Wt/Wt^* and *Mlkl^Wt/D139V^* mice after treatment with 150mg/kg 5FU or saline. Means ± SEM from one experiment in which three mice were sampled at each time point for each treatment group, similar results were obtained in an independent cohort. (J) Bone marrow cells (2×10^6^) from *Mlkl^Wt/Wt^* or *Mlkl^Wt/D139V^* mice on a CD45^Ly5.2^ background were mixed with 2×10^5^ wild type CD45^Ly5.1^ competitor bone marrow cells and transplanted into irradiated CD45^Ly5.1/Ly5.2^ recipients. Peripheral blood mononuclear cells from recipient mice were analysed after 56 days and then again at 180 days. Host contribution (CD45^Ly5.1/Ly5.2^) is depicted in gray, competitor (CD45^Ly5.1^) in white, and test (CD45^Ly5.2^) in black. The mean and standard error of the mean (SEM) are shown for 3 donors per genotype and 3-5 recipients per donor. (K) 2×10^6^ fetal liver cells (CD45^Ly5.2^; *Mlkl^Wt/Wt^*, *Mlkl^Wt/D139V^* or *Mlk^D139V/D139V^*) were transplanted into lethally irradiated recipients (CD45^Ly5.1/Ly5.2^) together with 2×10^5^ competitor bone marrow cells (CD45^Ly5.1^). Contribution to peripheral blood mononuclear cells was assessed 28 days after transplantation, and again at 180 days. Host contribution (CD45^Ly5.1/Ly5.2^) is depicted in gray, competitor (CD45^Ly5.1^) in white, and test (CD45^Ly5.2^) in black. Mean ± SEM are shown (2-10 donors per genotype, 2-6 recipients per donor).

### Homozygous *Mlkl^D139V^* fibroblasts are less sensitive to necroptotic stimuli and have low levels of MLKL protein

To examine if the constitutive activity of exogenously expressed MLKL^D139V^ results in an enhanced propensity for necroptosis in cells that express MLKL^D139V^ under the control of its endogenous promoter, we immortalized MDFs from *Mlkl^Wt/Wt^*, *Mlkl^Wt/D139V^* and *Mlkl^D139V/D139V^* littermates and from *Mlkl^−/−^* E19.5 pups. As expected, we observed no significant difference in the sensitivity of these cells to an apoptotic stimulus such as TNF plus Smac mimetic (Fig. 4A). However we observed a significant and consistent decrease in sensitivity to TNF induced necroptosis using three different pan-caspase inhibitors Q-VD-OPh, Z-VAD-fmk and IDUN-6556 in a *Mlkl^D139V^* dose dependent manner (Fig. 4A). While MDFs isolated from *Mlkl^D139V/D139V^* homozygotes were up to 60% less sensitive to TNF-induced necroptosis compared to *Mlkl^Wt/Wt^* MDFs, they were not completely resistant like *Mlkl^−/−^* MDFs (Fig. 4A).

**Figure 4:**
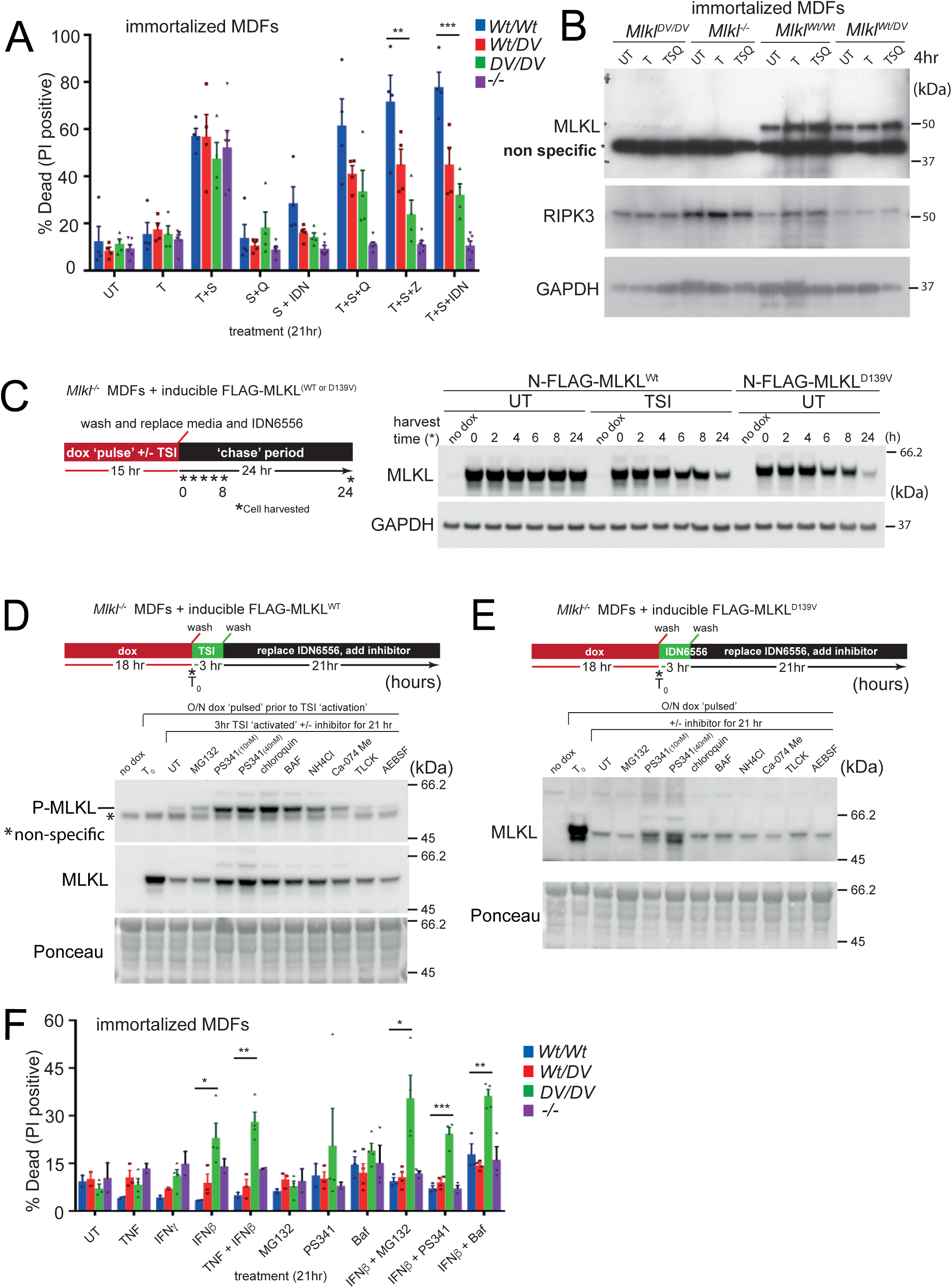
MLKL^D139V^ and activated MLKL^WT^ are cleared from cells via a mechanism that requires proteasome function and lysosomal acidification. MDFs were isolated from *Mlkl^Wt/Wt^*, *Mlkl^Wt/D139V^*, *Mlk^D139V/D139V^* or *Mlkl^−/−^ pups*, immortalized and stimulated as indicated for 21 hrs for quantification of PI-positive cells using flow cytometry (A), or for 4 hrs for western blot analysis (B). *Mlkl^−/−^* MDFs were stably transduced with doxycycline-inducible FLAG-MLKL^WT^ and FLAG-MLKL^D139V^ constructs to examine MLKL protein stability after doxycycline withdrawal (C) and in the presence of indicated compounds (D) and (E). (F) Immortalized MDFs from (A) stimulated as indicated for 21 hrs for quantification of PI-positive cells using flow cytometry. (A) and (F) represent mean ± SEM of 2-6 independent experiments. B-E are representative images of at least 3 independent experiments.

Surprisingly, while there were no obvious differences in the levels of MLKL^WT^ and MLKL^D139V^ protein following inducible exogenous expression (Fig. 1F), MLKL was virtually undetectable by Western blot in *Mlkl^D139V/D139V^* cells (Fig. 4B). There was, however, no significant reduction in *Mlkl* transcript levels in these cells suggesting that this reduction was post-transcriptionally regulated (Supp. Fig. 4A). The reduction in MLKL^D139V^ protein levels was also evident in whole body protein lysates prepared from E14 embryos (Supp. Fig. 4B). Lysates from E14 embryos also clearly show that *Mlkl^Wt/D139V^* heterozygotes have intermediate levels of MLKL, reflecting the intermediate sensitivity of *Mlkl^Wt/D139V^* MDFs to necroptotic stimuli (Fig. 4A).

### MLKL^D139V^ and RIPK3-phosphorylated wildtype MLKL is turned over in a proteasome and lysosome dependent manner

Measuring the half-life of endogenously expressed MLKL^D139V^ is not possible using conventional ‘pulse chase’ methods because this mutant protein induces necroptotic cell death, so we capitalized on our previous observation that an N-terminally FLAG-tagged MLKL 4HB forms a high molecular weight membrane-associated complex just like the untagged form, but, unlike the untagged version, does not kill cells (Hildebrand et al., 2014). Consistent with this observation, N-FLAG full-length mouse MLKL was phosphorylated by RIPK3 following stimulation with TSI, and formed a high molecular weight membrane associated complex, but did not induce cell death when inducibly expressed in *Mlkl^−/−^* MDFs (data not shown).

Using this system we were able to measure the half-life of MLKL by inducing N-FLAG-MLKL^WT^ or N-FLAG-MLKL^D139V^ expression in *Mlkl^−/−^* MDFs for 15 hours in doxycycline then washing and culturing them in the absence of doxycycline for a further 2-24 hours. In the absence of a stimulus (UT), the levels of N-FLAG-MLKL^WT^ remained consistent over the 24-hour period (Fig. 4C), indicating that wild type MLKL is a stable protein in MDFs. However, when these cells were treated with a necroptotic stimulus (TSI) the levels of wild type MLKL rapidly declined even though these cells were unable to undergo a necroptotic cell death. This indicates that RIPK3 induced phosphorylation, oligomerization or translocation to the membrane induces turnover of MLKL in a cell death independent manner. Consistent with the fact that untagged MLKL^D139V^ behaves as an auto-activated form of MLKL (Fig. 1E), the half-life of N-FLAG-MLKL^D139V^(4-6 hours) was similar to the WT version stimulated with TSI (Fig. 4C). Thus, the absence of endogenously expressed MLKL^D139V^ in E14 embryo lysates and cultured fibroblasts can be attributed to the reduced post-translational stability of this mutant auto-activated form of the protein.

To determine which cellular mechanism(s) are required for the clearance of activated MLKL, we included a series of proteasome, lysosome and specific protease inhibitors during the ‘chase’ period after doxycycline was withdrawn (schematic in Fig. 4D). The doses of these inhibitors were carefully titrated to minimize apoptotic cell death during the assay (Supp. Fig. 4C). Nevertheless, even at the very low doses used, the proteasome inhibitor PS341 reduced the clearance of TSI stimulated N-FLAG-MLKL^WT^ (Fig. 4D). This protection was particularly evident when specifically probing for phospho(p)-MLKL. Chloroquine, Bafilomycin and NH_4_Cl also partially protected against p-MLKL clearance (Fig. 4D). These agents have multifaceted actions, but interfere with the processes of lysosomal acidification and/or the fusion of autophagosomes/endosomes with lysosomes and thus prevent protein degradation by lysosomal proteases. Loss of total N-FLAG-MLKL^D139V^ was also prevented by PS341, however it was not possible to probe for p-MLKL as this activated form of MLKL is not phosphorylated in this assay due to the absence of TSI stimulation (Fig. 4E).

The reduced half-life of activated MLKL supports recent findings by others that mechanisms exist for the clearance of activated forms of MLKL (Gong et al., 2017; Yoon et al., 2017; Zargarian et al., 2017). Based on these findings we hypothesized that this MLKL-clearance mechanism limits the capacity of MLKL^D139V^ to kill *Mlkl^D139V^* hetero and homozygote cells in culture and *in vivo* by maintaining protein levels below a critical threshold. To test whether this protective mechanism could be overwhelmed, we incubated MDFs with agents that have been shown to induce *Mlkl* expression (TNF, interferons (IFN) β and γ) (Rodriguez et al., 2016; Rusinova et al., 2013; Tanzer et al., 2017; Thapa et al., 2013), or inhibit its turnover (proteasome and lysosome inhibitors). MLKL^D139V^ protein in untreated *Mlkl^D139V/D139V^* MDFs was undetectable by Western blot but became faintly detectable following stimulation with such stimuli (Fig. 4B & Supp. Fig. 4D). This correlates with moderate but statistically significant increases in cell death (particularly when compared with the lack of sensitivity to conventional necroptotic stimuli (Fig. 4A)), when exposed to IFNβ alone and in combination with proteasome or lysosome inhibitors (Fig. 4F). An allele-dose dependent sensitivity is also evident in primary MDFs (Supp. Fig. 4E). Together, these experiments provide evidence for the existence of steady-state MLKL surveillance and turn-over mechanisms that suppress cell death by lowering the abundance of activated MLKL below a killer threshold – both at the cellular *and* whole animal level.

Interestingly, genetic deletion of *Tnfr1, Myd88* and *Ifnar* did not provide any extension to the lifespan of *Mlkl^D139V^* homozygote pups (Table I), indicating that the removal of any one of these routes to NF-κB- and interferon-mediated gene upregulation is not sufficient to protect against a double allelic dose of *Mlkl^D139V^*. Similarly, combined genetic deletion of *Casp8* and *Ripk3* did not rescue or extend the life of *Mlkl^D139V/D139V^* mice, indicating that post-natal death is not mediated by bystander extrinsic apoptotic cell death that may occur secondary to initial waves of MLKL^D139V^-mediated necroptosis and associated inflammatory cytokine release (Table I). To test whether the death of *Mlkl^D139V/D139V^* neonates was mediated by activation of the inflammasome we also crossed this line with the *Caspase 1/11* null mouse strain (Kuida et al., 1995; Li et al., 1995). This did not enhance the lifespan of *Mlkl^D139V/D139V^* pups (Table I).

**Table I.** Postnatal lethality in *Mlkl^D139V^* homozygotes is independent of *Tnfr1*, *Myd88*, *Ripk3*, *Casp8*, *Casp1* and *Casp11*.

| *Mikl*<sup>Wt/D139V</sup> x *Mikl*<sup>Wt/D139V</sup> |  |  |  |  |  |  |  |  |  |
| --- | --- | --- | --- | --- | --- | --- | --- | --- | --- |
| stage genotyped | E14 | E18 | postnatal day 21 |  |  |  |  |  |  |
| C57BL/6 genetic background | Wt | Wt | Wt | *Tnfr1*<sup>-/-</sup> | *Myd88*<sup>-/-</sup> | *Ripk3*<sup>-/-</sup>, *C8*<sup>+/-</sup> | *Ripk3*<sup>-/-</sup>, *C8*<sup>-/-</sup> | *Ifnar*<sup>-/-</sup> | *C1*<sup>-/-</sup>, *C11*<sup>-/-</sup> |
| *Mikl*<sup>Wt/Wt</sup> | 58 (39) | 7 (10) | 15 (11) | 19 (15) | 3 (2) | 10(6) | 2(2) | 15 (11) | 1(1) |
| *Mikl*<sup>Wt/D139V</sup> | 70 (78) | 17 (20) | 30 (22) | 41 (30) | 6 (4) | 14(12) | 5(4) | 30 (22) | 2(2) |
| *Mikl*<sup>D139V/D139V</sup> | 28 (39) | 13 (10) | 0 (11) | 0 (15) | 0 (2) | 0 (6) | 0(2) | 0 (11) | 0(1) |
| total # genotyped | 156 | 37 | 45 | 60 | 9 | 24 | 7 | 45 | 3 |

( ) number of pups expected from mendellian segregation, calculated from total number of pups that genotyped, rounded to nearest whole number.

### Three of the four most frequent missense gene variants in human *MLKL* encode amino acid substitutions within or immediately adjacent to the brace region

Given the severe inflammatory phenotype of murine *Mlkl^D139V/D139V^* neonates and the significant defects in stress hematopoiesis observed in murine *Mlkl^Wt/D139V^* adults, we explored the prevalence of brace region variation in human *MLKL*. Examination of the gnomAD database (Lek et al., 2016), which contains human *MLKL* exome or genome sequence data from a total of over 141,456 individuals revealed that the second and third highest frequency human *MLKL* missense coding variants; rs34515646 (R146Q) and rs35589326 (S132P), alter the same brace helix (Table II, Fig. 5A). The 4^th^ most common human *MLKL* polymorphism, rs144526386 (G202*V) is a missense polymorphism identified exclusively in the context of a shorter splice isoform of MLKL (*) named ‘MLKL2’ (Arnez et al., 2015) (Table II, Fig. 5B). The full length canonical transcript of *MLKL* encodes a 471 amino acid protein, while alternatively spliced *MLKL2* encodes an isoform of MLKL that is 263 amino acids long and is missing a large portion of the pseudokinase domain which functions to repress the killing potential of the 4HB domain (Cai et al., 2014; Chen et al., 2014; Dondelinger et al., 2014; Hildebrand et al., 2014) and recruit co-effectors like RIPK3 and HSP90 ((Jacobsen et al., 2016; Petrie et al., 2018). Glycine202* is encoded by an extension to exon 9 that is unique to the *MLKL2* splice isoform (Fig. 5A, B).

**Table II.** Human *MLKL* brace helix polymorphism frequency

| Feature | human MLKLSNP |  |  |
| --- | --- | --- | --- |
|  | *R146Q- rs34515646* | *S132P- rs35589326* | *G202\*V- rs144526386* |
| CADD Score (phred-scaled) | 0.407 | 6.381 | 3.825 |
| UK Biobank- Total MAF (n) | 0.0253 (487,658) | 0.0161 (487,625) | 0.0147 (487,488) |
| gnomAD- Total MAF (n) | 0.0152 (141,339) | 0.0138 (141,442) | 0.01228 (141,400) |
| gnomAD- Highest MAF (n) population | 0.0252 (64,541)<br>European (Non-Finnish) | 0.0311 (5,185)<br>Ashkenazi Jewish | 0.0245 (5,184)<br>Ashkenazi Jewish |
| 1000 genomes- Total MAF (n) | 0.0052 (2,504) | 0.0088 (2,504) | 0.0102 (2,504) |
| 1000 genomes- Highest MAF (n) population | 0.018 (503)<br>European | 0.024 (489)<br>South Asian | 0.021(503)<br>European |

*n* - number of individuals sequenced MAF - Minor Allele Frequency -count \*alternate transcript

**Figure 5.**
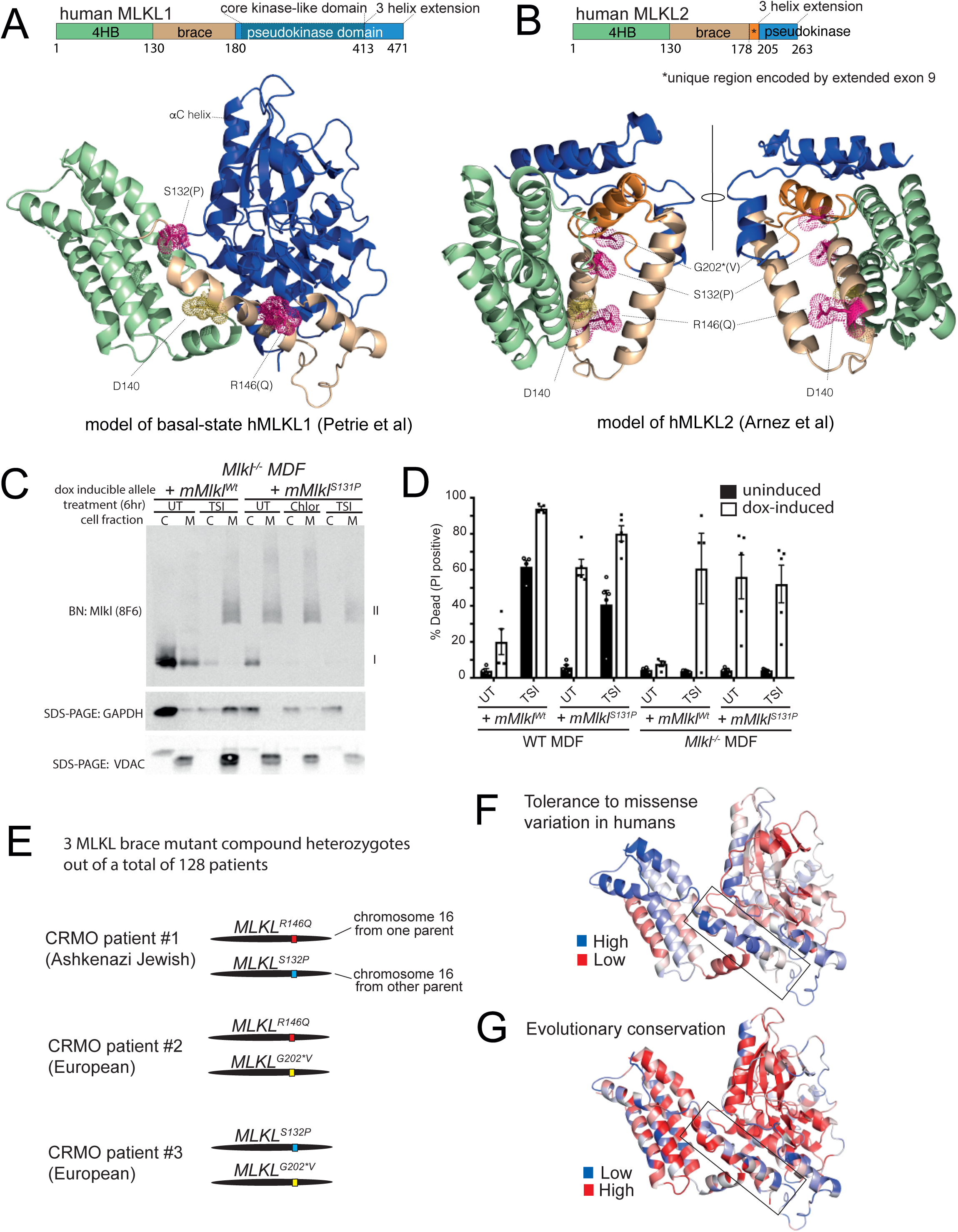
Three of the four highest frequency missense human MLKL SNPs encode non – conservative amino acid substitutions within or adjacent to the brace helix region. (A) S132 and R146 (magenta) are located on either side of D140 (yellow – equivalent to mouse D139) in the first human MLKL brace helix. Alternate amino acids encoded by human polymorphisms indicated in parentheses. (B) G202 is predicted to be on an α helix unique to MLKL splice-isoform 2 and to form an interface along with S132 and R146. The mouse equivalent of human rs35589326 (hMLKL^S132P^), mMLKL^S131P^, spontaneously forms membrane-associated high molecular weight complexes following Blue Native (BN) PAGE (C) and kills MDFs (D) in the absence of extrinsic necroptotic stimuli when expressed in mouse dermal fibroblasts for 6 (C) and 21 hrs respectively (D). C; cytoplasmic fraction, M; crude membrane fraction, TSI; TNF, Smac-mimetic and IDN6556. (E) Schematic showing brace helix variant combinations identified as alleles *in trans* in 3 CRMO patients. (F) MTRs are mapped onto the structure of MLKL to show regions that have low tolerance to missense variation in the human population (red) and regions that have increased tolerance to missense variation (blue), normalized to the gene’s MTR distribution. (G) Evolutionary conservation Multiple sequence alignment (MSA) conservation scores are mapped onto the structure of MLKL to show regions that are highly conserved through evolution (red) and regions that are less conserved through evolution (blue). (C) is representative of 2 independent experiments, (D) mean ± SEM of 4-5 independent experiments.

While the amino acid substitution *MLKL^R146Q^* is classified as ‘tolerated’ and ‘benign’ by SIFT/POLYPHEN 2 algorithms (Adzhubei et al., 2013; Sim et al., 2012) (Supp. Table IV), R146 of human MLKL shows NMR chemical shift perturbations in the presence of the negatively charged phospholipids IP3 and IP6, indicating a possible role in membrane association and disruption (Dovey et al., 2018; Quarato et al., 2016). Ser-132 lies at the intersection of a dynamic disordered loop and the first structured residue of the conserved brace helix 1 (Fig. 5A) (Murphy et al., 2013; Petrie et al., 2018; Su et al., 2014). A Serine-to-Proline substitution at this position is predicted to significantly impact the conformation of the immediately adjacent W133 (brace helix) and in turn, the closely situated W109 (4 helix bundle) (Supp. Fig. 5A). When mapped to a model of MLKL splice-isoform 2 (Arnez et al., 2015) Glycine 202* is predicted to be on an isoform 2-specific helix and to form an interface along with S132 and R146 of brace helix 1. While the precise structural consequence of these three brace polymorphisms is unknown, modelling of human MLKL predicts that disruption in the brace region favours adoption of an activated conformation (Petrie et al, 2018). Consistent with this prediction, the murine equivalent of the human S132P variant, mMLKL^S131P^, formed high molecular weight membrane-associated complexes and killed MDFs in the absence of a necroptotic stimulus (Fig. 5 C, D) when expressed at close to endogenous levels (Supp. Fig. 5B).

### MLKL brace helix variants appear *in trans* at a higher frequency in a cohort of CRMO patients than in healthy controls

To investigate if human MLKL brace region polymorphisms play a role in human autoinflammatory disease we examined their frequency in cohorts suffering from Ankylosing Spondylitis (AS), chronic recurrent multifocal osteomyelitis (CRMO), Guillain Barre Syndrome (GBS) and Synovitis, Acne, Pustulosis, Hyperostosis and Osteitis (SAPHO) Syndrome. The individual minor allele frequencies of R146Q, S132P and G*202V are not enriched in these disease cohorts relative to healthy controls when population distribution is accounted for (Supp. Tables IV and V). However these alleles occur *in trans* (making ‘compound heterozygotes’ – schematic in Fig. 5E) in 3 out of 128 CRMO patients. This is 29 times the frequency that these combinations are observed in healthy NIH 1000 genomes samples (where there are only 2 compound heterozygotes for these polymorphisms out of 2504 healthy individuals sequenced), or at 10-12 times the frequency when only European CRMO patients and two separate healthy European control populations were compared (Table III).

**Table III.** Human MLKL brace helix compound heterozygotes in CRMO vs Healthy Controls

| Population | Frequency of relevant compound Hets |  |  | 2 tailed p value |  |
| --- | --- | --- | --- | --- | --- |
|  | CRMO (n) | Healthy controls (n) | CRMO:Healthy<sup>a</sup> | Fisher's exact | χ square with Yates |
| Global | 0.023 (3/128) | 0.0008 (2/2504*)<br>NIH 1KG | 29:1 | 0.001 | 0.0001 |
| European | 0.02 (2/101) | 0.002 (1/503*)<br>NIH 1KG | 10:1 | 0.074 | 0.1215 |
| European | 0.02 (2/101) | 0.0017 (25/14,542*)<br>QUT controls | 12:1 | n/a | 0.0022 |

Frequency of CRMO patients that are compound heterozygotes (see Supp. Fig 5c for schematic)
*n* - number of individuals sequenced

## DISCUSSION

In contrast to apoptosis, necroptosis is widely held to be an inflammatory form of cell death. However, definitive evidence for this proposition has yet to emerge. Because MLKL is activated by inflammatory stimuli such as TNF it is very difficult to separate cause from effect. The identification of an auto-activating mutant of MLKL (*Mlkl^D139V^*) in mice has allowed us to explore the consequences of inappropriate necroptosis in the absence of such confounding factors. Furthermore it has led to significant insights into developmental processes sensitive to MLKL activation and into physiological mechanisms that exist to neutralize activated MLKL. These turnover mechanisms critically control cell fate, determining whether auto-active MLKL^D139V^ is present at a sufficient level to promote cell death.

While MLKL phosphorylation might serve as an immuno-histochemical marker for necroptosis ordinarily, in the *Mlkl^D139V^* mice it is not possible to pinpoint exactly which cell type/s undergo necroptosis. Nevertheless, the presence of high levels of circulating pro-inflammatory cytokines in *Mlkl^D139V/D139V^* pups at E19.5 relative to *Mlkl^Wt/Wt^* and *Mlkl^Wt/D139V^* littermates suggests that necroptosis and ensuing inflammation occurs in the sterile *in utero* environment. This is not enough to overtly retard prenatal development or affect hematopoietic cell populations other than by moderately reducing circulating platelet levels. However, upon birth and/or exposure to the outside environment the capacity of homozygous *Mlkl^D139V/D139V^* pups to suppress *MLKL^D139 V^*activity appears overwhelmed and they die within days of birth. This is clearly a dose-dependent effect because both *Mlkl^D139V/Wt^* and *Mlkl^D139V/null^* heterozygous mice are viable. We therefore speculate that transcriptional upregulation of *Mlkl^D139V^* overwhelms the turnover and/or membrane repair mechanisms that counteract MLKL activation (Gong et al., 2017; Yoon et al., 2017). Post-natal death cannot be prevented by combined deficiencies in *Ripk3* and *Casp8* or indeed deficiency of any other inflammatory gene that we tested, including *Tnfr1, Myd88* or *Ifnar*. This further supports the idea that excessive MLKL-induced necroptosis can generate an inflammatory response in the absence of other inflammatory mediators. Difficulty with suckling due to inflammatory infiltration of the head and neck and resulting failure to thrive is one possible explanation for the lethality. However, the narrow window of mortality for *Mlkl^D139V/D139V^* pups and marked pericardial immune infiltration make heart failure another potential cause of sudden neonatal death.

One of the most unexpected findings from our study is the physiological importance of endogenous mechanisms that limit the ability of activated MLKL to kill cells. While others have recently shown that an ESCRT dependent repair mechanism can help protect membranes from limited MLKL damage it was not feasible to demonstrate the physiological relevance of this finding (Gong et al., 2017; Yoon et al., 2017). Our data suggest both proteasomal and lysosomal mechanisms also exist to dispose of activated MLKL. While proteasomal degradation is usually considered to be cytoplasmic and completely separate from lysosomal degradation, it was notable that low doses of either the proteasome inhibitor, PS341, or chloroquine (that inhibits lysosome acidification) limited p-MLKL degradation to very similar extents. This creates the possibility that these mechanisms or the previously described ESCRT mechanism intersect. Finally, the ability of these mechanisms to hold heterozygous levels of active MLKL in check without deleterious consequences *in vivo* supports the idea that direct inhibition of activated MLKL may be an effective means to therapeutically prevent unwanted necroptotic cell death.

The *Mlkl^D139V^* brace mutant mouse strain may be a useful model to study the role of necroptosis in human health and disease. According to current allele frequencies in gnomAD, up to 8% of individuals globally are heterozygous for missense *MLKL* gene variants within the brace-coding region. This percentage of people with brace variants indicates that this region is highly tolerant to missense mutation (Fig. 5F, Supp. Fig. 5C). High tolerance to missense variation in a coding sequence is often used to filter out potential pathogenic variants in human genetic studies because it indicates that such variations are likely to be functionally neutral (Traynelis et al., 2017). However, the first brace helix is both highly evolutionarily conserved yet also tolerant of missense mutations in the human population (Fig. 1D, Fig. 5F,G). Furthermore *in vivo* and *in vitro* data show that amino acid substitutions in the brace region have profound effects on MLKL function (Davies et al., 2018; Quarato et al., 2016). Therefore, overlayed with structural, biochemical, cell and animal-based evidence of function, it is tempting to speculate that these human MLKL brace region variants have accumulated not simply by chance, but through positive evolutionary selection. While defective emergency hematopoiesis is likely to be subject to *negative* evolutionary selection, *Mlkl^D139V^* mouse-derived HSCs are only defective following chemo- or radio-ablation. Given that these forms of HSC depletion are unlikely to have been a significant selective force during human evolution, we speculate that these human brace polymorphisms have achieved high frequencies in the human population because they have conferred a selective advantage to infectious disease. Evidence for positive selection has been found for over 300 immune-related gene loci and many of these have been found to be associated with the incidence of autoimmune and autoinflammatory disease in modern humans (Gutierrez-Arcelus et al., 2016; Ramos et al., 2015). Many of these variants have also been mechanistically linked to defense against a particular pathogen (Karlsson et al., 2014; Ramos et al., 2015). While increased numbers and examination of independent cohorts will be required to confirm the statistical enrichment of human MLKL brace variants occurring in *trans* in CRMO, this patient cohort offers a tantalizing first clue into their potential as modifiers of complex/polygenic inflammatory disease.

## EXPERIMENTAL PROCEDURES

### Mice

All mice were backcrossed to C57BL/6 mice for >10 generations or generated on a C57BL/6J background. *Mlkl*^−*/-*^, *Tnfr1^−/−^, Myd88^−/−^, IFNAR1^−/−^, Ripk3^−/−^, Casp8^−/−^* and *Casp1/Casp11^−/−^ mice* were generated as described (Adachi et al., 1998; Beisner et al., 2005; Hwang et al., 1995; Kuida et al., 1995; Li et al., 1995; Murphy et al., 2013; Newton et al., 2004; Peschon et al., 1998). Mice designated as E19.5 were obtained by Caesarean section from mothers that received progesterone injections at E17.5 and E18.5. An independent mouse strain that carried the D139V mutation in the *Mlkl* gene (MLKL^D139V^ CRISPR) was generated using CRISPR/Cas9 as previously described (Wang et al., 2013). Briefly, one sgRNA of the sequence GGAAGATCGACAGGATGCAG (10ng/μl), an oligo donor of the sequence ATTGGAATACCGTTTCAGATGTCAGCCAGCCAGCATCCTGGCAGCAGGAAGATCGACAGGTTGCAGAAGAAGACGGgtgagtctcccaaagactgggaaagagtaggccagggttgggggtagggtgg (10ng/uL) and Cas9 mRNA (5ng/μL) were injected into the cytosol of C57BL/6J zygotes. Mice were sequenced across the mutated region to confirm incorporation of the altered codon and analysis was performed after at least 2 back-crosses to C57BL/6. The relevant Animal Ethics Committee approved all experiments.

### Linkage analysis

We mapped the chromosomal location of the *Plt15* mutation by mating affected mice to 129/Sv *Mpl^−/−^* mice to produce N_2_ (backcross) and F_2_ (intercross) generations. A genome wide scan using 20 N_2_ mice with the highest platelet counts (287±74×10^6^/ml, compared with 133±75×10^6^/ml for the overall population, Fig. 1A) localized the mutation to a region of chromosome 8 between *D8Mit242* and *D8Mit139* and linkage to this region was then refined. Analysis of the F_2_ population revealed a significant reduction in the frequency of mice homozygous for *C57BL/6* alleles in this interval (e.g. *D8Mit200* 3/81 F_2_ mice homozygous *C57BL/6*, p=2.2×10^−5^ χ^2^-test), suggesting the *Plt15* mutation results in recessive lethality. The refined 2.01 Mb interval contained 31 annotated genes, only five of which appeared to be expressed both in the hematopoietic system and during embryogenesis (http://biogps.gnf.org/): *Dead box proteins 19a* and *19b* (*Ddx19a* and *Ddx19b*), *Ring finger and WD repeat domain 3* (*Rfwd3*), *Mixed lineage kinase domain like* (*Mlkl*), and *WD40 repeat domain 59* (*Wdr59*). Sequencing identified a single mutation, an A to T transversion in *Mlkl* that was heterozygous in all mice with an elevated platelet count.

### Reagents

#### Antibodies

Rat-anti mRIPK3 and rat anti-mMLKL 8F6 (selected for affinity to residues 1-30 of mouse MLKL) and rat anti-MLKL 3H1 (MLKL brace region) were produced in-house. Anti-Pro Caspase 8 (#4927) and GAPDH (#2113) were purchased from Cell Signaling Technology. Anti-mouse P-MLKL (ab196436) and anti-Actin (ab5694) were purchased from Abcam. Anti-VDAC (AB10527) was purchased from Millipore. FC-hTNF was produced in house and used at a final concentration of 100ng/mL. Recombinant mouse IFNγ and β were purchased from R&D Systems (Minneapolis, MN, USA) Q-VD-OPh and ZVAD were purchased from MP Biomedicals (Seven Hills, NSW, Australia). Smac mimetic also known as Compound A, and the caspase inhibitor IDN-6556 were a gift from TetraLogic (Malvern, PA, USA). Propidium iodide, doxycycline, and bafilomycin were purchased from Sigma-Aldrich (Castle Hill, NSW, Australia).

### Cell line generation and culture

Primary mouse dermal fibroblasts were prepared from skin taken from the head and body of E19.5 pups delivered by C-section or from the tails of adult mice as described (Etemadi et al., 2013). Primary MDFs were immortalized by stable lentiviral transduction with SV40 large T antigen. Immortalized MDFs were stably transduced with exogenous mouse and human MLKL cloned into the pFTRE 3G vector, which was generated by Toru Okamoto, and allows doxycycline-inducible expression as described (Murphy et al., 2013). Cells were maintained in culture as previously described (Tanzer et al., 2017).

### Cell death assays

Cell death assays were performed as described previously using 5 × 10^4^ MDFs per well in 24 well tissue culture plates (Murphy et al., 2013). Doxycycline (20 ng/mL) was added together with death stimuli. Fc-hTNF was produced in house and used at 100ng/mL, Compound A Smac mimetic and IDN6556 were used at 500 nM and 5 μM respectively. ZVAD and QVD-OpH were used at 25 μM and 10 μM respectively. Mouse and human interferons gamma and beta were used at 30 ng/mL, PS341 and MG132 at 2 nM and 200 nM respectively and Bafilomycin at 300 nM.

### MLKL turn-over assays

5 × 10^4^ MDFs per well were plated in 24 well tissue culture plates and allowed to settle. Doxycycline (20 ng/mL) +/-TNF, Smac Mimetic and IDN6556 was added. After 15 hr, ‘no dox’ and ‘0’ wells were harvested. Media was removed from remaining wells and cells were washed with PBS and fresh media containing IDN6556 was re-added. Wells were then harvested 2, 4, 6, 8 and 24 hours from this point. Cells were harvested by direct lysis in reducing SDS-PAGE lysis buffer.

### MLKL protection assays

5 × 10^4^ MDFs per well were plated in 24 well tissue culture plates and allowed to settle. Doxycycline (20 ng/mL) was added. After 18 hrs, ‘no dox’ and ‘T_0_’ samples were harvested. Media was removed and cells washed before addition of fresh media containing TSI or IDN alone for 3 hrs. Cells were washed again and media restored with IDN6556 alone (UT), or IDN6556 + inhibitor (MG132 (200 nM), PS341 (10-40 nM), Choloroquin (50 μM), Bafilomycin (300 nM), Ca-074 Me (20 μM), TLCK (100 μM) and AEBSF (100 μM)) for a further 21 hours. Cells were harvested by direct lysis in reducing SDS-PAGE lysis buffer.

### Transmission Electron Microscopy

Murine dermal fibroblasts prepared from mice of the indicated genotypes were untreated or stimulated with the indicated agents for the indicated hours. Then, cells were fixed with 2% glutaraldehyde in 0.1 M phosphate buffer, pH 7.4, postfixed with 2% OsO_4_, dehydrated in ethanol, and embedded in Epok 812 (Okenshoji Co.). Ultrathin sections were cut with an ultramicrotome (ultracut N or UC6: Leica), stained with uranyl acetate and lead citrate, and examined with a JEOL JEM-1400 electron microscope. The viability of a portion of these cells was determined by measuring LDH release as described previously (Murai et al., 2018).

### Mouse histopathology

Caesarian-sectioned E19.5 and Day P2/3 pups were euthanized by decapitation and fixed in 10% buffered formalin. 5 μm coronal sections were taken at 200 μm intervals for the full thickness of the head, 5 μm sagittal sections were taken at 300 μm intervals for the full thickness of the body. A thorough examination of these sections was performed by histopathologists Aira Nuguid and Tina Cardamome at the Australian Phenomics Network, Melbourne. Findings were confirmed by Veterinary Pathologist Prof. John W. Finney, SA Pathology, Adelaide and clinical Pathologist Prof. Catriona McLean, Alfred Hospital, Melbourne.

### Measurement of relative thymic cortical thickness

Representative images of thymus sections were analysed to determine relative cortical thickness using ImageJ. Briefly, medullary areas were identified on the basis of H and E staining and removed from the larger thymus structure using the Image J Image Calculator function to isolate the cortical region. The thickness of the cortical region, defined by the radius of the largest disk that can fit at a pixel position, was determined using the Local Thickness plugin in ImageJ (http://www.optinav.info/Local_Thickness.htm).

### Immunohistochemistry

Following terminal blood collection, P0 and P3 pups were fixed for at least 24 hours in 10% buffered formalin and paraffin embedded before microtomy. Immunohistochemical detection of Cleaved caspase 3 (Cell Signaling Technology #9661) and CD45 (BD) was performed as described previously (Rickard et al., 2014b).

### Cytokine quantification

All Plasma was stored at −80°C prior to cytokine analyses. Cytokines were measured by Bioplex Pro mouse cytokine 23-plex assay (Bio- Rad #M60009RDPD) according to manufacturer’s instructions. When samples were designated ‘<OOR’ (below reference range) for a particular cytokine, they were assigned the lowest value recorded for that cohort (as opposed to complete exclusion or inclusion as ‘zero’ which would artificially inflate or conflate group averages respectively). Values are plotted as fold change relative to the mean value for the *Wt/Wt* samples, and p values were calculated in Microsoft Excel using a 2 tailed TTEST, assuming unequal variance. Data is only shown for cytokines that displayed statistically significant differences between genotypes at either of or both day E19.5 and day P3.

### Hematological Analysis

Blood was collected from P0 and P3 pups into EDTA coated tubes using heparinized glass capillary tubes from the neck cavity immediately after decapitation. After centrifugation at 500G for 5 min, 5-15 μL of plasma was carefully removed and this volume was replaced with PBS. Blood cells were resuspended and diluted between 8-20 fold in DPBS for automated blood cell quantification using an ADVIA 2120 hematological analyzer within 6 hours of harvest. Blood was collected from adult mice retro-orbitally into tubes containing EDTA and analyzed using an ADVIA120 automated hematological analyzer (Bayer).

### Transplantation Studies

Donor bone marrow or fetal liver cells were injected intravenously into recipient *C57BL/6-CD45^Ly5.1/Ly5.2^* mice following 11Gy of gamma-irradiation split over two equal doses. Recipient mice received neomycin (2 mg/mL) in the drinking water for 4 weeks. Long term capacity of stem cells was assessed by flow cytometric analysis of donor contribution to recipient mouse peripheral blood and/or hematological organs up to 6 months following engraftment. Recovery from cytotoxic insult was assessed by automated peripheral blood analysis at regular times following treatment of mice with 150 mg/kg 5-fluorouracil (5-FU).

### Flow Cytometry

To analyze the contribution of donor and competitor cells in transplanted recipients, blood cells were incubated with a combination of the following antibodies: Ly5.1-PE, Ly5.2-FITC, Ly5.2-biotin or Ly5.2 PerCPCy5.5 (antibodies from Becton Dickenson, Ca). If necessary, cells were incubated with a streptavidin PECy5.5 (BD), mixed with propidium iodide (Sigma) and analysed on a LSRI (BD Biosciences) flow cytometer. To analyse the stem- and progenitor cell compartment, bone marrow cells were incubated with biotinylated or Alexa700 conjugated antibodies against the lineage markers CD2, CD3, CD4, CD8, B220, CD19, Gr-1 and Ter-119.

For stem and progenitor cell detection antibodies against cKit, Sca-1, CD48, AnnexinV, CD105, FcγRII/III or CD135 in different combinations (see antibody list for details). Finally FluoroGold (AAT Bioquest Cat#17514) was added for dead cell detection. Cells were then analysed on LSRII or Fortessa1 (BD Biosciences) flow cytometers.

### Reactive Oxygen Species (ROS) detection

ROS was detected by using Chloromethyl-H_2_DCFDA dye according to the manufacturer’s instructions (Invitrogen Cat#C6827). In brief, bone marrow cells were loaded with 1μM Chloromethyl-H_2_DCFDA for 30 minutes at 37°C. Loading buffer was then removed, and cells were placed into 37°C StemPro-34 serum free medium (ThermoFisher Cat#10639011) for a 15 minute chase period. After incubation cells were placed on ice and stained with surface antibodies suitable for FACS analysis. Cells were analysed using a LSRII flow cytometer (Becton Dickinson).

### Quantitative PCR

RNA was prepared using Trizol (Invitrogen) according to the manufacturer’s instructions and 10μg was used for first strand cDNA synthesis using SuperScript II (Life Technologies). ∼0.5 μg of cDNA was then used in a TaqMan PCR reaction with Universal PCR mastermix and murine Mlkl (Mm1244222_n1) and GAPDH (Mm99999915_m1) Taqman probes (ThermoFisher) on an ABI 7900 Fast Real-Time PCR instrument (Applied Biosystems). Mlkl expression relative to GAPDH control was determined using SDS version 2.3 program (Applied Biosystems) and expressed as ΔCT values.

### Statistics (Mouse and cell-based assays)

Please consult figure legends for description of error bars used. All P values were calculated in Microsoft Excel or Prism using an unpaired, two tailed t-test, assuming unequal variance. * p ≤ 0.05, ** p ≤ 0.01, ***p ≤ 0.005

### Whole Exome Sequencing

DNA from CRMO probands and their family members (when available) was purified from saliva or blood and prepared for whole exome sequencing (WES). The samples underwent WES at several different times, enriched using the Agilent SureSelect Human All Exon V4, V5 or V6+UTR (Agilent Technologies) before sequencing at either Otogenetics, Inc (Atlanta, GA), Beckman Coulter Genomics (Danvers, MA), or at the University of Iowa Genomics Core (Iowa City, IA). The fastq files were quality-checked and processed to vcf format as described previously (Cox et al., 2017). Variants for all samples were called together using GATK’s Haplotype Caller (McKenna et al., 2010) and were recalibrated and hard-filtered in GATK as described previously (Cox et al., 2017). Variants were annotated with minor allele frequencies (MAFs) from 1000 genomes (Genomes Project et al., 2015), ExAC and gnomAD (Lek et al., 2016) and with information regarding the effect of each variant using SNPSift/SNPEff (Cingolani et al., 2012). The databases used for annotation were dbNSFP2.9 (Liu et al., 2016a) (for MAFs) and GRCh37.75 for protein effect prediction.

### Ancestry Determination

Ancestry was determined for each CRMO proband using the LASER software package (Wang et al., 2014). A vcf file including ten probands at a time was uploaded to the LASER server and the TRACE analysis was selected using the Worldwide panel. For probands with indeterminate ancestry using the Worldwide panel, the European and Asian panels were used. Principal component values for each proband were plotted using R Statistical Software and the code provided in the LASER package.

### MLKL variant quantification

#### 1000 Genomes

Vcf files from 1000 genomes were annotated and filtered as described previously (Cox, 2018). Values for MLKL variants rs35589326 (S132P), rs34515646 (R146Q), and rs144526386 (G202V) as well as all MLKL coding variants were queried and tabulated for allele and genotype count for participants of all ancestry (n=2504), and for those of European ancestry (n=503). Compound heterozygous variants were evident due to the phasing of all variants in the 1000 genomes dataset. *CRMO:* Allele and genotype counts for all MLKL coding variants were tabulated in probands of European ancestry (n=101) and for all probands (n=128). Compound heterozygous variants were identified using parental sequence data. *AS:* DNA from all subjects in AS cohort were genotyped using the Illumina CoreExome chip following standard protocols at the Australian Translational Genomics Centre, Princess Alexandra Hospital, Brisbane. Bead intensity data was processed and normalized for each sample and genotypes called using the Illumina Genome Studio software. All the samples listed in the table have been passed quality control process. *GB:* Genotyping was performed in an ISO15189-accredited clinical genomics facility, Australian Translational Genomics Centre (ATGC), Queensland University of Technology. All samples were genotyped by Illumina HumanOmniExpress (OmniExpress) BeadChip (Blum et al., 2018). *QUT controls:* A collection of healthy control data of verified European ancestry from various cohort studies, complied by the Translational Genomics Group, QUT and typed on an Illumina CoreExome microarray. Includes data from the The UK Household Longitudinal Study, led by the Institute for Social and Economic Research at the University of Essex and funded by the Economic and Social Research Council. The survey was conducted by NatCen and the genome-wide scan data were analysed and deposited by the Wellcome Trust Sanger Institute. Information on how to access the data can be found on the understanding Society website https://www.understandingsociety.ac.uk/.

### Statistical Analysis (Human data)

Statistical comparisons were performed at the level of allele frequency or the level of compound heterozygote sample frequency using either a Fisher’s exact test or a Chi-Squared test with Yates correction as specified under each table. Compound heterozygous variants were quantified and compared at the individual rather than the allelic level, where individuals with and without qualifying variants were compared at the allelic level.

### Web resources

gnomAD – https://gnomad.broadinstitute.org/ http://asia.ensembl.org

OrthoDB – https://www.orthodb.org

CADD – https://cadd.gs.washington.edu/

Clustal Omega – https://www.ebi.ac.uk/Tools/msa/clustalo/

WEBLOGO – https://weblogo.berkeley.edu/logo.cgi

Missense Tolerance Ratio (MTR) Gene Viewer – http://biosig.unimelb.edu.au/mtr-viewer

UK biobank – https://www.ukbiobank.ac.uk

## AUTHOR CONTRIBUTIONS

Conceptualization: JMH, MK, JMM, WSA, JS.

Methodology: JMH, JMM, WA, JS, HA, JR

Investigation: JMH, MK, IJM, ZL, AC, SM, EJP, MAS, MCT, SNY, CH, SEG, JC, MDS, PG, ECJ, KR, AT, JS, TSS, JGZ, CCA, GMT, ECH, TAW, DS, CAG, JC, AH, NS, SKS, DC, DM, MSH, CGV, CM, MB, SLM, JMM

Resources: JMH, JGZ, VA, RML, AGB, BWD, MAFS, NV, DH, MK, WZ, KW, NV, JT, SB, JR, CGV, PM, MAB, BTK, PJF, JMM, WSA, JS

Writing-Original Draft: JMH, WA, JS

Writing – Review and Editing: JMH, MK, EJP, PAC, PM, PJF, SLM, JMM, WSA, JS Supervision-JMH, MP, PJF, HN, JMM, WA, JS

Funding Acquisition-JMH, JMM, WA, JS

## ACKNOWLEDGEMENTS

We thank all the following people for their technical assistance; Jiami Han, Cynthia Liu, Jasmine McManus, Janelle Lochland (WEHI). Aira Nuguid and Tina Cardamone (APN histopathology – The University of Melbourne). Thomas Boudier (WEHI Centre for Dynamic Imaging). The WEHI Histology Service, WEHI Antibody Facility and WEHI Bioservices. Y. Uchiyama and S. Kakuta who advised the interpretation of the results of TEM. Victoria Jackson and Annette Jacobsen for important insight and discussion. The generation of *Mlkl^D139V^* mice by CRISPR/Cas9 was performed by Andrew Kueh and Marco Herold (WEHI MAGEC laboratory) supported by the Australian Phenomics Network (APN) and the Australian Government through the National Collaborative Research Infrastructure Strategy (NCRIS) program.

This work is supported by; Project grant (1105023) and Fellowships (0541951 and 1142669) from the Australian National Health and Medical Research Council (NHMRC) to JMH. Project grant (1105023) and Fellowship (1107149) from the NHMRC to JS. Program grant (1113577) and Fellowship (1058344) from the NHMRC (WSA). JMM-Project grant (1124735) and Fellowship (1105754) from the NHMRC (JMM). NIH training grants T32GM008629 and T32GM082729-01 (AJC). R01AR059703 from the National Institute of Arthritis and Musculoskeletal and Skin Diseases (NIAMS) at the National Institutes of Health (PJF and AGB), the Marjorie K. Lamb Professorship PJF. Program grant (1113577) and Fellowship (1058344) from the NHMRC (WSA). Grants-in-Aid from Scientific Research (B) 17H04069 (to HN) from Japan Society for the Promotion of Science (JSPS), and Scientific Research on Innovative areas 26110003 (to HN), the Japan Agency for Medical Research and Development (AMED) through AMED-CREST with a grant number JP18gm1210002 (to HN), and Private University Research Branding project (to HN) from a MEXT (Ministry of Education, Culture, Sports, Science and Technology). Victorian International Research Scholarship (Z. Liu and MCT). Australia Postgraduate Award (CAD). SLM acknowledges funding from NHMRC grants (1144282,1142354 and 1099262), The Sylvia and Charles Viertel Foundation, HHMI-Wellcome International Research Scholarship and Glaxosmithkline. Fellowship from the Lorenzo and Pamela Galli Charitable Trust (ECJ). NHMRC grants 1107425 and 1045549 and The Sylvia & Charles Viertel Senior Medical Research Fellowship (MP). DBA was supported by the Jack Brockhoff Foundation (JBF 4186, 2016) and NHMRC Fellowship (APP1072476). Supported in part by the Victorian Government’s OIS Program. NHMRC IRIISS and Victorian Government Operational Infrastructure Support schemes. NHMRC Project and Targeted Research grants 1006769, 512672 and 512381 to MFS.

MAB acknowledges the Department of Industry, Innovation, Science, Research and Tertiary Education Collaborative Research Network and Diabetes Australia for their support. IJM was supported by the Victorian Cancer Agency, and by generous support from the Felton Bequest.

We gratefully acknowledge the contribution of genotype data by Dr Yorgi Mavros (University of Sydney), Professor Nick Martin (QIMR), Professor Jim Rosenbaum (Oregon Health and Science University), and Professor Maxime Breban and the Groupe Française d’Etude Génétique des Spondylarthrites (GFEGS). We are grateful to Professor BP Wordsworth of the University of Oxford, UK for access to genotype data on ankylosing spondylitis cases collected in studies funded, in part, by Arthritis Research UK (grants 19536 and 18797), by the Wellcome Trust (grant 076113) and by the Oxford Comprehensive Biomedical Research Centre ankylosing spondylitis chronic disease cohort (theme A91202).

## SUPPLEMENTARY FIGURE LEGENDS

**Supp. Fig. 1.**
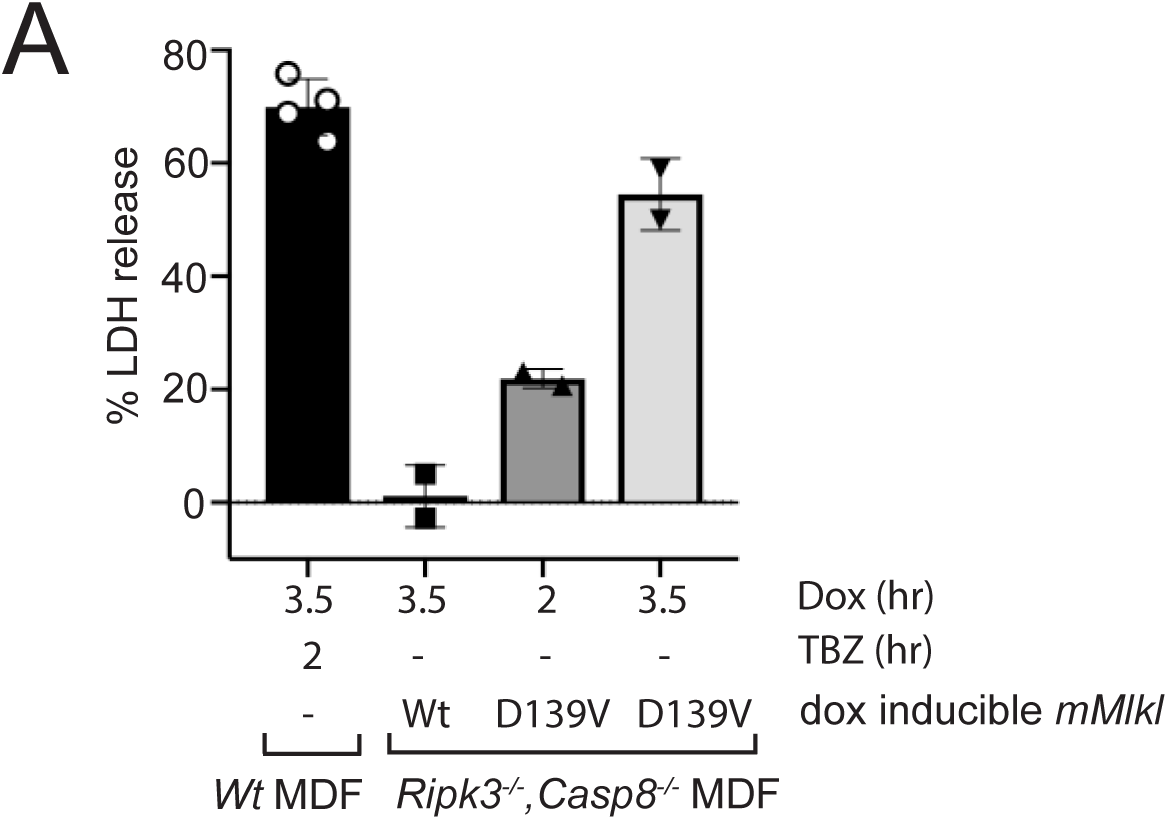
(A) Viability of non-transduced *Wt* MDFs or *RIPK3^−/−^, Caspase8^−/−^* MDFs expressing dox-inducible *Mlkl^Wt^* or *Mlkl^D139V^* was monitored by measuring LDH release at the indicated time points post addition of TNF (T), Birinpant (B) and ZVAD-fmk (Z) or doxycycline. These conditions correspond to those used for TEM analyses (Fig. 1G).

**Supp. Fig. 2.**
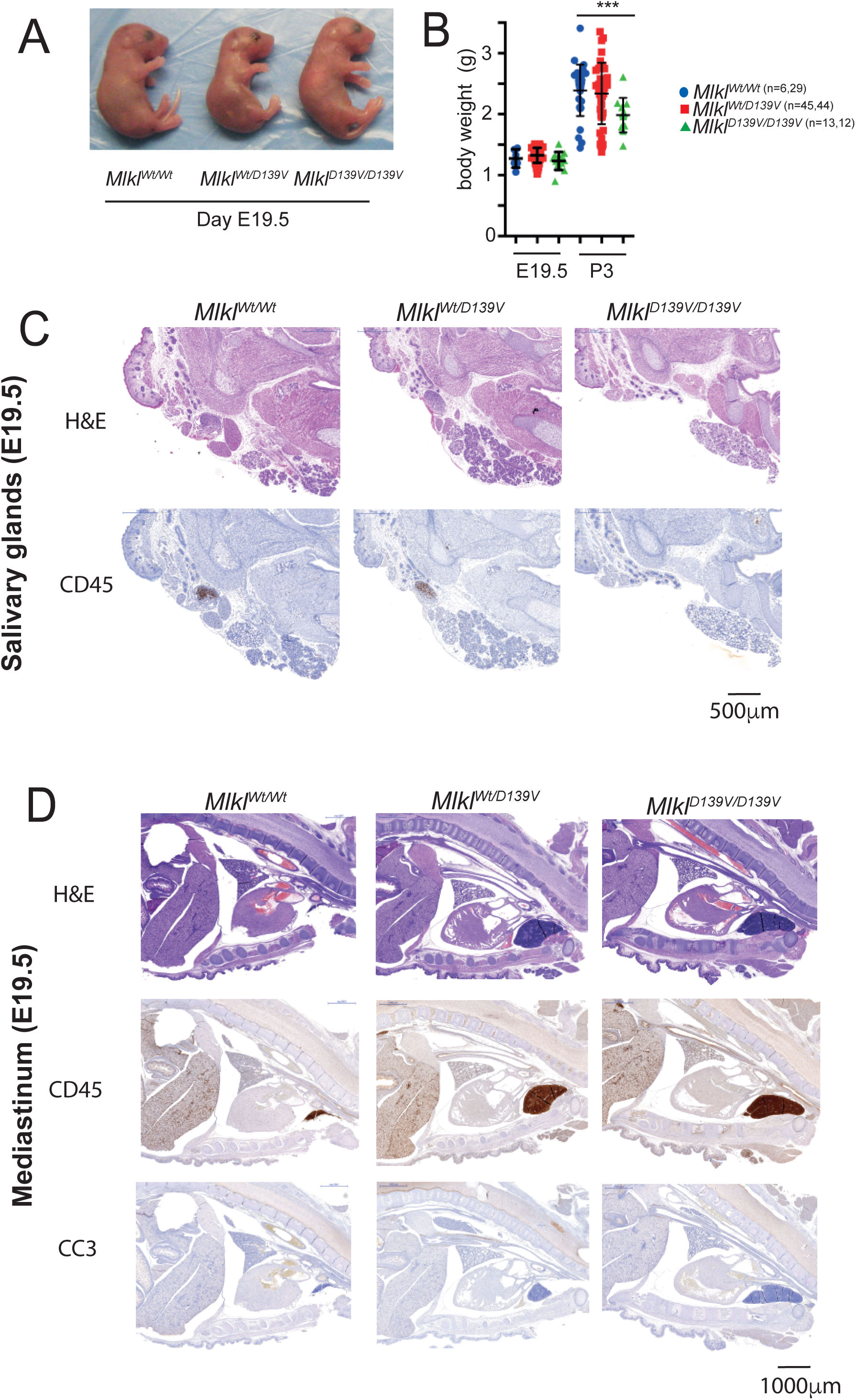
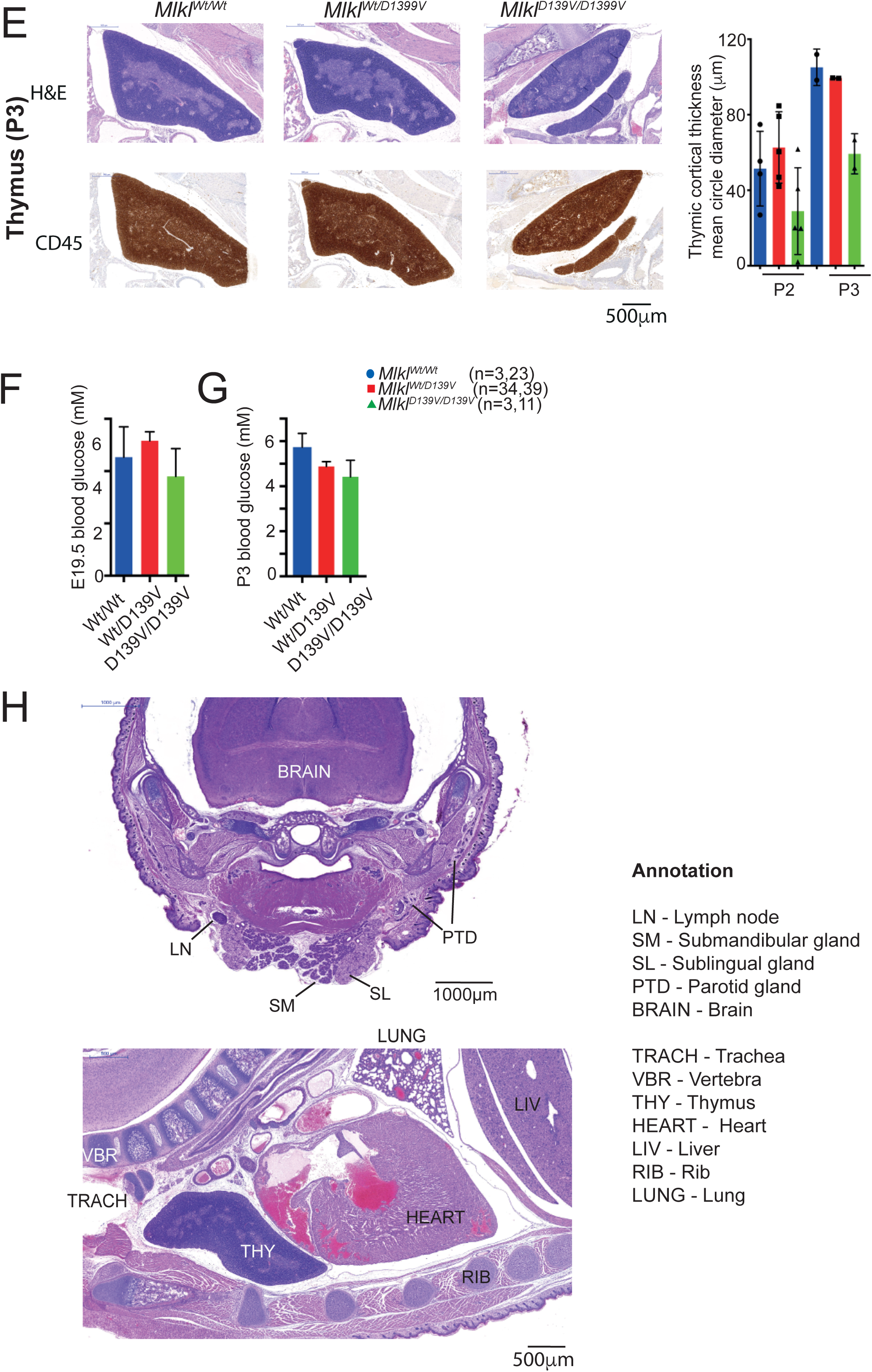
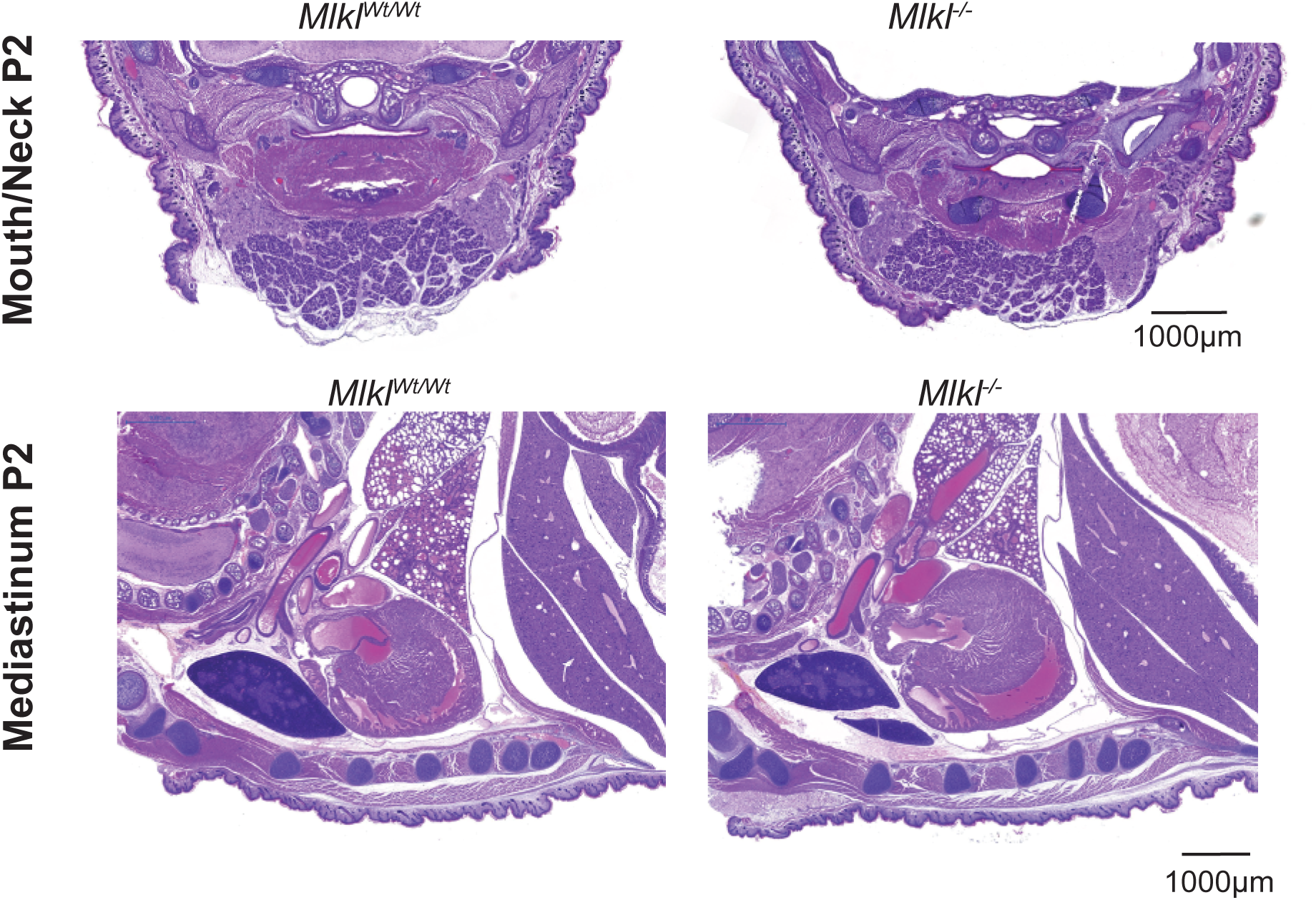
(A) Macroscopic appearance of E19.5 pups of indicated genotypes after Caesarean delivery. (B) Body weights of *Mlkl^Wt/Wt^*, *Mlkl^Wt/D139V^*, *Mlk^D139V/D139V^* mice at E19.5 and postnatal Day 3. (C) Serial mandible sections from E19.5 pups stained with H&E and anti-CD45. (D) H&E and anti-CD45 or cleaved caspase-3 (CC3) stained section of E19.5 mediastinum. (E) Serial sections of thymi from postnatal day 3 pups stained with H&E and anti-CD45 and quantification of thymic cortical thickness. (F-G) Blood glucose measured at E19.5 and postnatal day 3 (non-fasting) plotted as mean ± SEM for n=3-39 pups per genotype. (H) Anatomical annotation of head and mediastinum of postnatal day 2 *Mlkl^Wt/Wt^* pup. (I) Coronal section of postnatal day 2 pup mouth/neck region and mediastina stained with H&E.

**Supp. Fig. 3.**
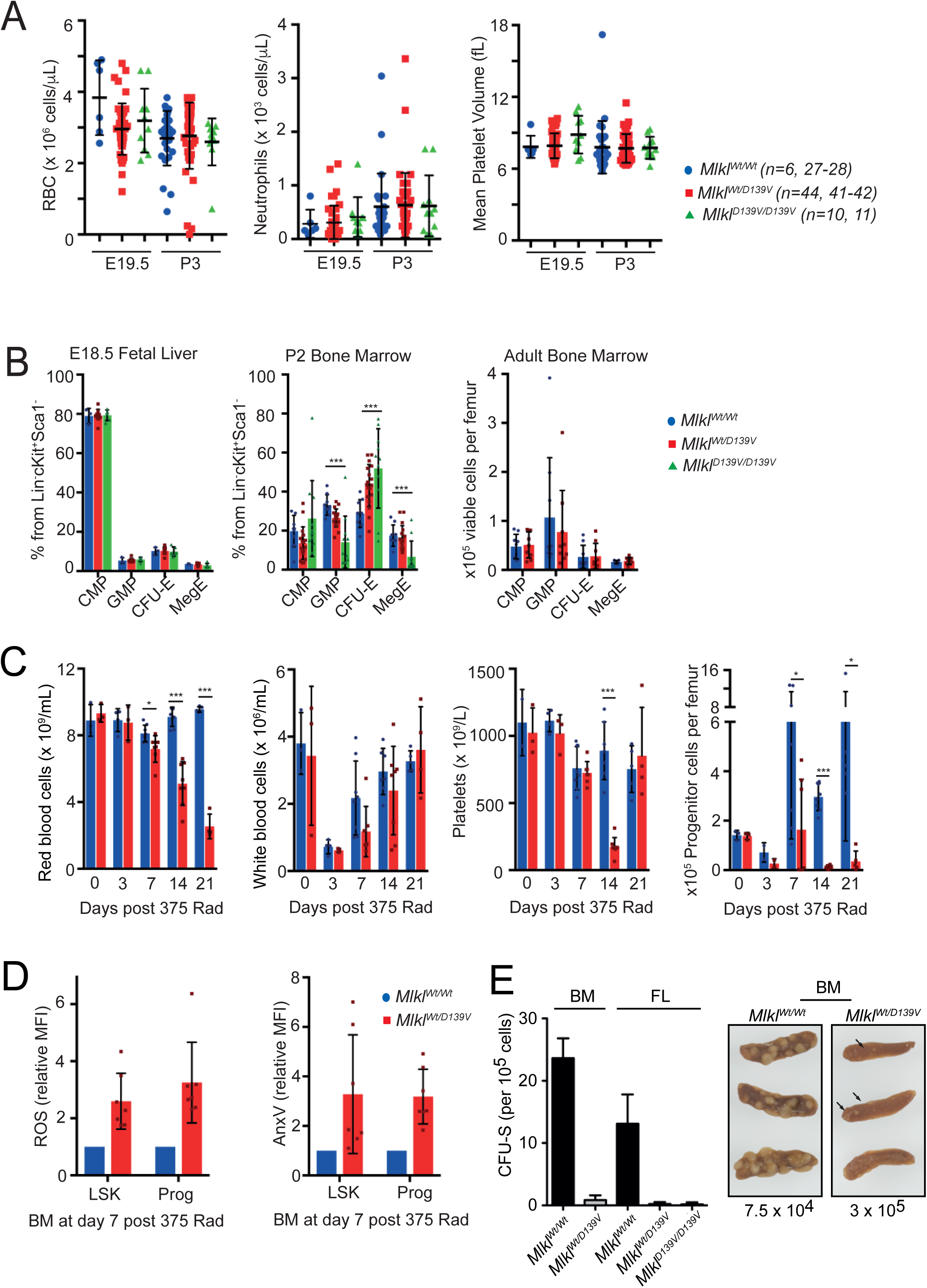
(A) Numbers of red blood cells (RBC), neutrophils and mean platelet volume in the peripheral blood of E19.5 and P3 pups. (B) Common myeloid progenitors (CMP, Lineage^−^IL7Rα^−^Sca1^−^cKit^+^CD34^+^ FcγRII/III^−^), granulocyte-macrophage progenitors (GMP, Lineage^−^IL7Rα^−^cKit^+^Sca1^−^CD150^−^Endoglin^−^FcγRII/II^+^), Colony-forming units-erythroid (CFU-E, Lineage^−^IL7Rα^−^cKit^+^Sca1^−^ CD150^−^FcγRII/III^−^Endoglin^hi^), and megakaryocyte-erythroid progenitors (MegE, Lineage-IL7Rα^−^Sca1^−^cKit^+^CD150^+^Endoglin^low^ FcγRII/III^−^) in E18.5 fetal livers, P2 bone marrow cells and adult bone marrow, presented as a percent from Lin-cKit+Sca1-cell fractions. Mean ± SD, n=3-6 (E18.5), n=9-11 (P2 BM), n=9 (adult BM) per genotype. (C) Recovery of red blood cells, white blood cells, platelets and bone marrow progenitor cells (Lineage-Sca^−^Kit^+^) in *Mlkl^Wt/Wt^* and *Mlkl^Wt/D139V^* mice following 375 Rad whole body irradiation. (D) Relative amount of ROS and AnnexinV in LSK and progenitor cells was determined 7 days post irradiation. (E) Bone marrow (BM) or fetal liver (FL) cells (7.5×10^4^ −3×10^5^) from mice of the indicated genotypes (*Mlkl^Wt/Wt^*, *Mlkl^Wt/D139V^* or *Mlk^D139V/D139V^*) were transplanted into lethally irradiated recipients and spleens were removed for enumeration of CFU-S after 8 days. Mean ± SEM from 2-8 donors. Spleens taken from recipients of *Mlkl^Wt/Wt^* or *Mlkl^Wt/D139V^* bone marrow (7.5 ×10^4^ or 3.0 ×10^5^ cells transplanted respectively) were photographed to detail the size and number of colonies. *Mlkl^Plt15/+^* cells generated very small colonies at low frequency (arrows).

**Supp. Fig. 4.**
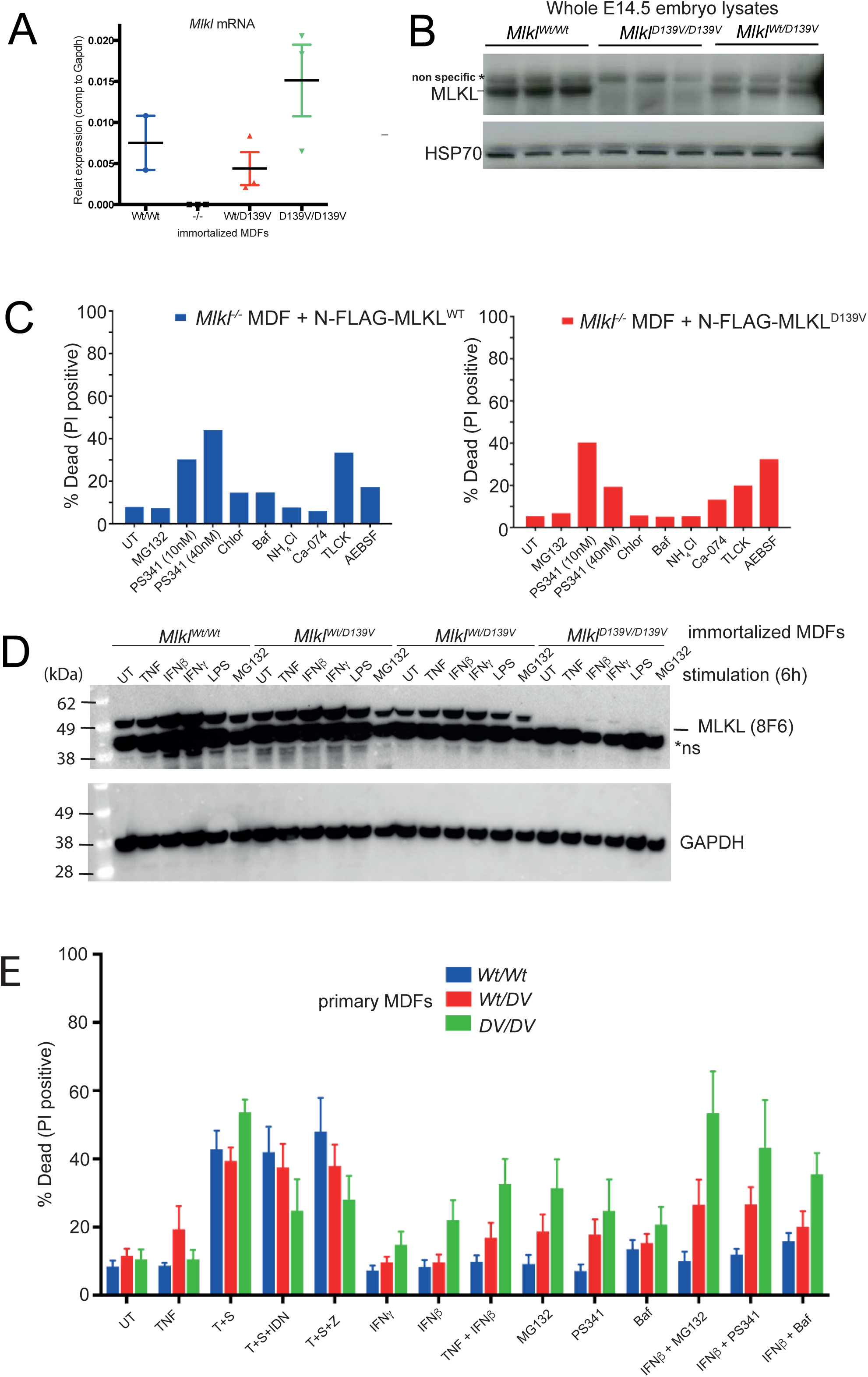
(A) Mouse *Mlkl* mRNA levels quantified using TaqMan probes. (B) E14.5 whole embryo lysates from 3 pups per genotype were probed by western blot for relative MLKL protein levels. (C) Viability of cells following 21 hr incubation with inhibitors used in Fig 4D-E. Representative of 3 similar experiments. (D) MDFs were treated as indicated for 21 hours. Whole cell lysates were analysed by western blot for levels of MLKL. (E) Primary MDFs were isolated from *Mlkl^Wt/Wt^*, *Mlkl^Wt/D139V^* or *Mlk^D139V/D139V^* mice and stimulated as indicated for 21hrs for quantification of PI positive cells using flow cytometry.

**Supp. Fig. 5.**
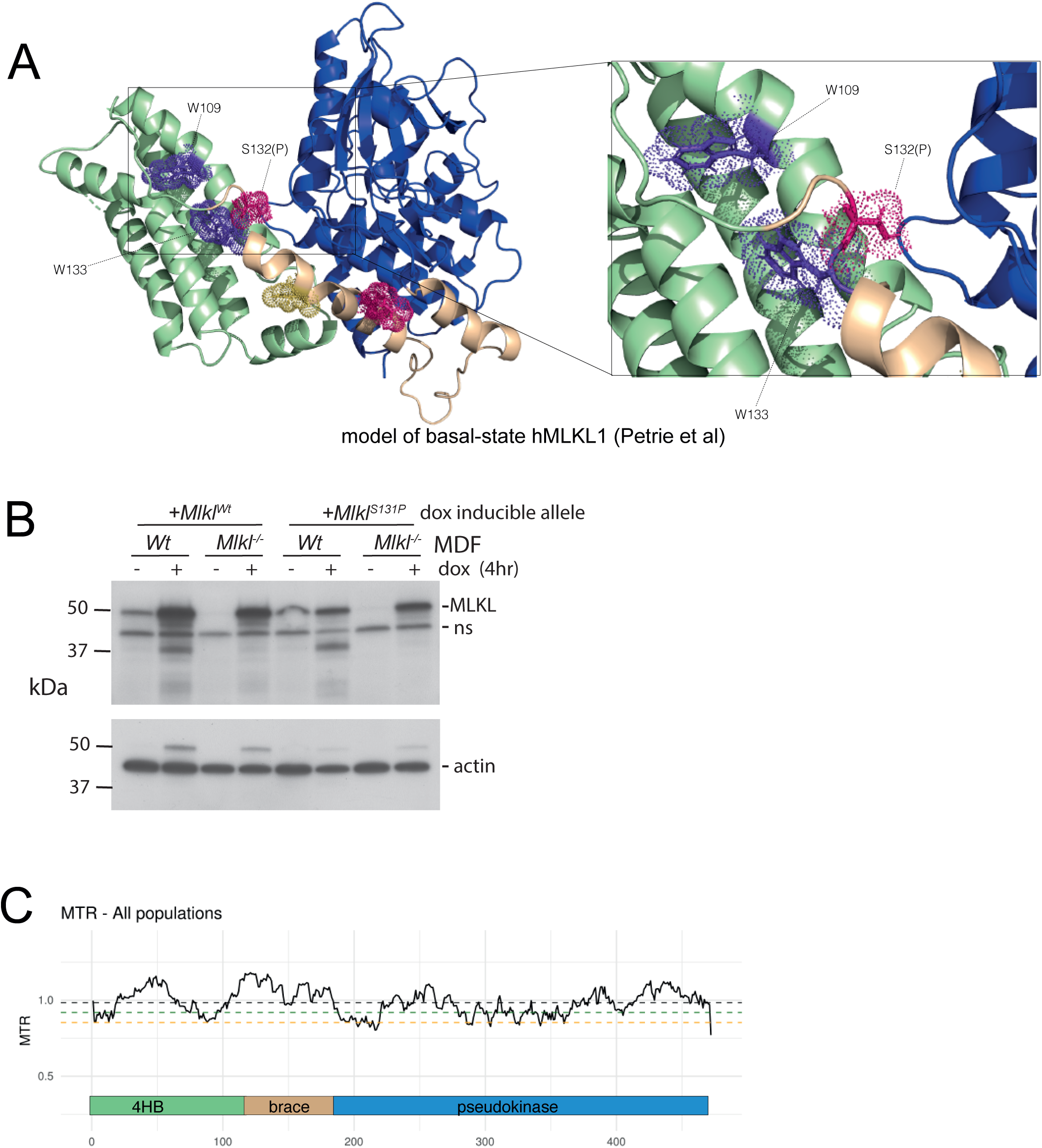
(A) A proline in position 132 of human MLKL is predicted to significantly impact the conformation of the immediately adjacent W133 (brace helix) and in turn, the closely situated W109 (4 helix bundle). (B) MDFs stably transduced with doxycycline inducible constructs expressing mouse MLKL^S131P^ were analysed by western blot for MLKL levels after 4 hrs dox induction. (C) Missense Tolerance Ratio (MTR) distribution for human MLKL using gnomAD exome data.

**Table SI.**
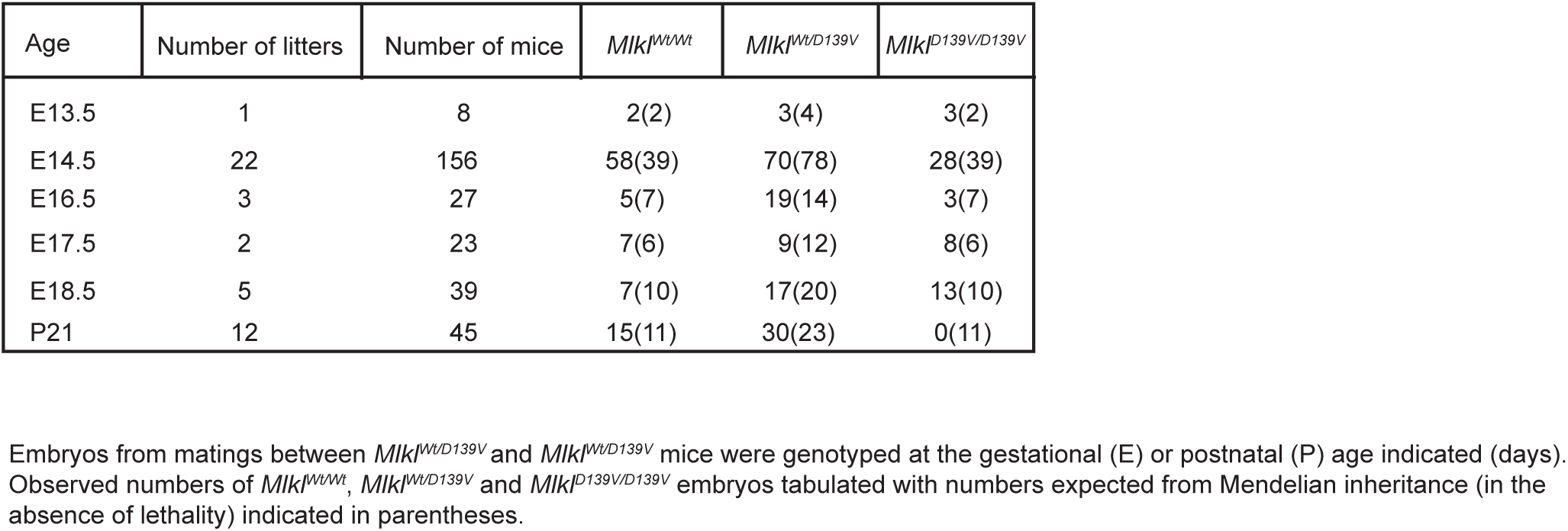
Outcome of *Mlkl^Wt/D139^* ^x^ *Mlkl^Wt/D139^* cross

| Age | Number of litters | Number of mice | *Mikl<sup>Wt/Wt</sup>* | *Mikl<sup>Wt/D139V</sup>* | *Mikl<sup>D139V/D139V</sup>* |
| --- | --- | --- | --- | --- | --- |
| E13.5 | 1 | 8 | 2(2) | 3(4) | 3(2) |
| E14.5 | 22 | 156 | 58(39) | 70(78) | 28(39) |
| E16.5 | 3 | 27 | 5(7) | 19(14) | 3(7) |
| E17.5 | 2 | 23 | 7(6) | 9(12) | 8(6) |
| E18.5 | 5 | 39 | 7(10) | 17(20) | 13(10) |
| P21 | 12 | 45 | 15(11) | 30(23) | 0(11) |

Embryos from matings between *Mikl<sup>Wt/D139V</sup>* and *Mikl<sup>Wt/D139V</sup>* mice were genotyped at the gestational (E) or postnatal (P) age indicated (days). Observed numbers of *Mikl<sup>Wt/Wt</sup>*, *Mikl<sup>Wt/D139V</sup>* and *Mikl<sup>D139V/D139V</sup>* embryos tabulated with numbers expected from Mendelian inheritance (in the absence of lethality) indicated in parentheses.

**Table SII.** Outcome of *Mlkl^Wt/D139 x^ Mlkl^null/null^* cross

| Age | Number of litters | Number of mice | *Mikl<sup>Wt/null</sup>* | *Mikl<sup>D139V/null</sup>* |
| --- | --- | --- | --- | --- |
| P21 | 8 | 40 | 19 (20) | 21 (20) |

Surviving progeny from matings between *Mikl<sup>null/null</sup>* and *Mikl<sup>Wt/D139V</sup>* mice were genotyped at postnatal day 21. Observed numbers are tabulated, with numbers expected from mendelian inheritance in the absence of lethality indicated in parentheses.

**Table SIII.** Outcome of CRISPR-*Mlkl^Wt/D139V^* ^x^ CRISPR-*Mlkl^Wt/D139V^* cross

| Age | Number of litters | Number of mice | *Mikl<sup>Wt/Wt</sup>* | *Mikl<sup>Wt/D139V</sup>* | *Mikl<sup>D139V/D139V</sup>* |
| --- | --- | --- | --- | --- | --- |
| P21 | 7 | 36 | 12 (9) | 24 (18) | 0 (9) |

Surviving progeny from matings between CRISPR-induced *Mikl<sup>Wt/D139V</sup>* mice were genotyped at postnatal day 21. Observed numbers are tabulated, with numbers expected from mendelian inheritance in the absence of lethality indicated in parentheses.

**Supp. Table IV.** Human *MLKL* brace helix variants – individual MAFs in CRMO vs Healthy Con-

| Feature | human *MLKL* SNP |  |  |
| --- | --- | --- | --- |
|  | *R146Q*- *rs34515646* | *S132P*- *rs35589326* | *G202\*V*- *rs144526386* |
| 1000 genomes- EU MAF (n) | 0.0179 (503) | 0.0149 (503) | 0.0209 (503) |
| U.Iowa CRMO cohort Total MAF (n) | 0.0273 (128) | 0.0234 (128) | 0.0234 (128) |
| U.Iowa CRMO cohort EU MAF (n) | 0.0347 (101) | 0.0198 (101) | 0.0198 (101) |
| U.Iowa CRMO TOTAL vs 1000 genomes Total MAF | p= 0.0009 | p= 0.0329 | p= 0.057 |
| U.Iowa CRMO EU vs 1000 genomes EU MAF | p= 0.1687 | p= 0.5423 | p= 0.99 |
| SIFT Score (classification) | MLKL1- 0.569 (TOLERATED)<br>MLKL2- 0.536 (TOLERATED) | MLKL1- 0.25 (TOLERATED)<br>MLKL2- 0 (DELETERIOUS) | MLKL2- 0.069 (TOLERATED) |
| POLYPHEN-2 Score (classification) | MLKL1- 0.114 (BENIGN) | MLKL1- 0.996 (PROBABLY DAMAGING) | n/a |

*n* - number of unrelated individuals sequenced
MAF - Minor Allele Frequency -count
\*alternate transcript
*p* - 2-tailed fisher's exact *p*-value by comparing the allele counts in cases and controls

**Supp. Table V.**
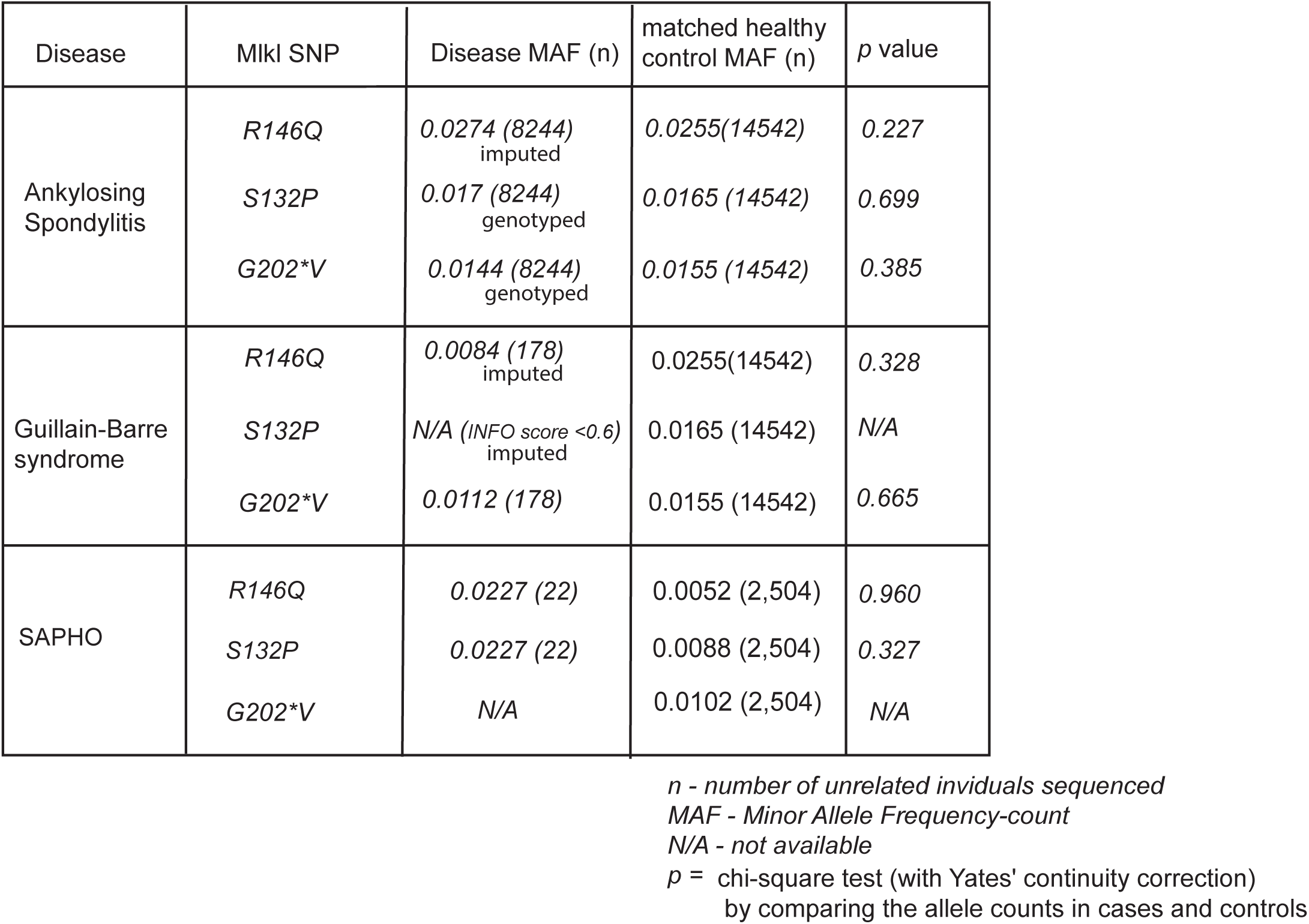
Human MLKL brace helix individual MAFs in AS, GB and SAPHO vs Healthy Controls

| Disease | MIKI SNP | Disease MAF (n) | matched healthy control MAF (n) | *p* value |
| --- | --- | --- | --- | --- |
| Ankylosing Spondylitis | *R146Q* | 0.0274 (8244)<br>imputed | 0.0255(14542) | 0.227 |
|  | *S132P* | 0.017 (8244)<br>genotyped | 0.0165 (14542) | 0.699 |
|  | *G202\*V* | 0.0144 (8244)<br>genotyped | 0.0155 (14542) | 0.385 |
| Guillain-Barre syndrome | *R146Q* | 0.0084 (178)<br>imputed | 0.0255(14542) | 0.328 |
|  | *S132P* | N/A (*INFO* score <0.6)<br>imputed | 0.0165 (14542) | N/A |
|  | *G202\*V* | 0.0112 (178) | 0.0155 (14542) | 0.665 |
| SAPHO | *R146Q* | 0.0227 (22) | 0.0052 (2,504) | 0.960 |
|  | *S132P* | 0.0227 (22) | 0.0088 (2,504) | 0.327 |
|  | *G202\*V* | N/A | 0.0102 (2,504) | N/A |

*n* - number of unrelated individuals sequenced
MAF - Minor Allele Frequency-count
N/A - not available
*p* = chi-square test (with Yates' continuity correction) by comparing the allele counts in cases and controls

